# Base Editing Rescue of Seizures and SUDEP in *SCN8A* Developmental Epileptic Encephalopathy

**DOI:** 10.1101/2025.04.09.647983

**Authors:** Caeley M. Reever, Alexis R. Boscia, Tyler C.J. Deutsch, Mansi P. Patel, Raquel M. Miralles, Shrinidhi Kittur, Erik J. Fleischel, Atum M. L. Buo, Matthew S. Yorek, Miriam H. Meisler, Charles R. Farber, Manoj K. Patel

## Abstract

*SCN8A* encodes the sodium channel Na_v_1.6; mutations in *SCN8A*, particularly gain-of-function variants, cause *SCN8A* developmental and epileptic encephalopathy (DEE), a severe epilepsy syndrome characterized by seizures, cognitive dysfunction, and seizure-induced death. The recurrent *SCN8A* variant R1872W impairs channel inactivation, causing neuronal hyperexcitability and seizures. Current treatments for *SCN8A* DEE are often ineffective, highlighting the need for targeted therapies. Here we employed base editing to correct the R1872W *SCN8A* variant. A modified adenine base editor (*SCN8A*-ABE) was packaged within dual PhP.eB-adeno-associated viruses (AAVs) and administered to R1872W mice at P2. *SCN8A*-ABE significantly increased survival and reduced or eliminated seizures. Neuronal hyperexcitability and heightened sodium currents were attenuated with *SCN8A*-ABE treatment, and behavioral comorbidities were improved. These effects were achieved by approximately 30% conversion of mutant to wildtype *SCN8A* transcripts. These findings demonstrate base editing as an effective targeted therapeutic approach for *SCN8A* DEE by addressing the underlying genetic cause.

## Introduction

*SCN8A* encodes the voltage-gated sodium channel Na_v_1.6, which is expressed primarily in the central and peripheral nervous systems and is essential for the initiation and propagation of action potentials in excitable cells^1^. It is highly expressed in both excitatory and inhibitory neurons and found concentrated at nodes of Ranvier and the axon initial segment (AIS)^2^. Pathogenic variants in *SCN8A* lead to developmental and epileptic encephalopathy (DEE)^3,4^. To date, over 500 individuals have been identified with pathogenic *SCN8A* variants, and many exhibit a gain-of-function (GOF) phenotype^5^. Patients experience refractory seizures and a range of comorbidities, including movement disorders, developmental delays, and cognitive dysfunction. The risk of sudden unexpected death in epilepsy (SUDEP) is high in the *SCN8A* DEE patient population^6^. A recurrent single nucleotide polymorphism (SNP) at residue R1872 accounts for approximately 10-15% of all *SCN8A* DEE cases and is associated with neonatal seizures^4,7,8^. The CGG codon for arginine, at residue 1872, contains a CpG dinucleotide, which is prone to a high mutation rate due to its vulnerability to cytosine deamination^9^. In Na_v_1.6 channels harboring the R1872W mutation, the loss of arginine in the C-terminal domain destabilizes its interaction with the inactivation gate, resulting in altered sodium channel inactivation and increased neuronal excitability that underlies the severe seizure phenotype in *SCN8A* DEE^4,10–12^. Current treatment options are limited to the use of anti-seizure medications (ASMs) that target sodium channels as a means of suppressing seizure activity without affecting the underlying genetic cause. Unfortunately, many patients are treatment resistant while others suffer intolerability issues and/or side effects associated with ASMs^13,14^.

Here, we have used adenine base editors (ABEs) to precisely revert the single-nucleotide change of adenine to guanine at the R1872W residue of *SCN8A*^15,16^, addressing the underlying cause of the disorder. We identified a highly effective construct and its paired editor for correction of R1872W, referred to as *SCN8A*-ABE. In vivo, *SCN8A*-ABE treatment increased survival and either significantly reduced seizure frequency or completely abolished seizure onset in a mouse model of *SCN8A* DEE. *SCN8A*-ABE treatment resulted in editing efficiencies observable in RNA transcripts of approximately 30%, reverting the mutant tryptophan (W; TGG) to wildtype arginine (R; CGG). No A to G editing was observed within 26 transcript sites, or on 2 protein coding genes, and all but 1 (2.4%) of 112 intronic/intergenic regions with potential for off target ABE effects. Electrophysiology recordings revealed attenuation of neuronal hyperexcitability and a suppression of aberrant sodium channel activity in ABE-treated mice. Additionally, we observed mitigation of multiple behavioral abnormalities associated with *SCN8A* DEE after ABE-treated mice. In summary, we demonstrate that base editing therapy rescues the disease phenotype of *SCN8A* DEE by targeting the underlying genetic cause and ameliorating key aspects of the disorder. These studies highlight the immense potential of base editing techniques as a novel therapeutic approach for not only *SCN8A* DEE, but other genetic epilepsies as well.

## Results

### Design and Validation of Adenine Base Editor Targeting SCN8A R1872W

The *SCN8A* variant R1872W (Fig. 1.a) results from a CGG to TGG substitution on the coding DNA strand. We targeted the negative (complementary) strand to enable an adenine to guanine conversion. This ultimately resulted in a TGG to CGG correction upon completion of the intermediate base editing conversion of adenine to inosine (Fig. 1.b)^16^. Additionally, there is no canonical NGG PAM, the typical recognition site for Cas9^17^, near the R1872W loci in a utilizable position for guide RNA (sgRNA) constructs. Therefore, we explored base editors with fluid PAM sequences in cell screens to identify the most efficient constructs. We designed a set of guide (sgRNA) sequences capable of utilizing 23 published ABE deaminase constructs with diverse PAM recognition profiles to revert R1872W. We chose to pair these into 16 testable constructs (Supp. Table 1) based on reported PAM preferences^18–21^ and high predictive accuracy from BE-HIVE^22^, CRISPOR^23^, and CRISPR RGEN tools^24^. Each construct was designed to target the same adenine at the R1872W *SCN8A* locus and took into consideration the available PAM sequences in the area.

**Figure 1:**
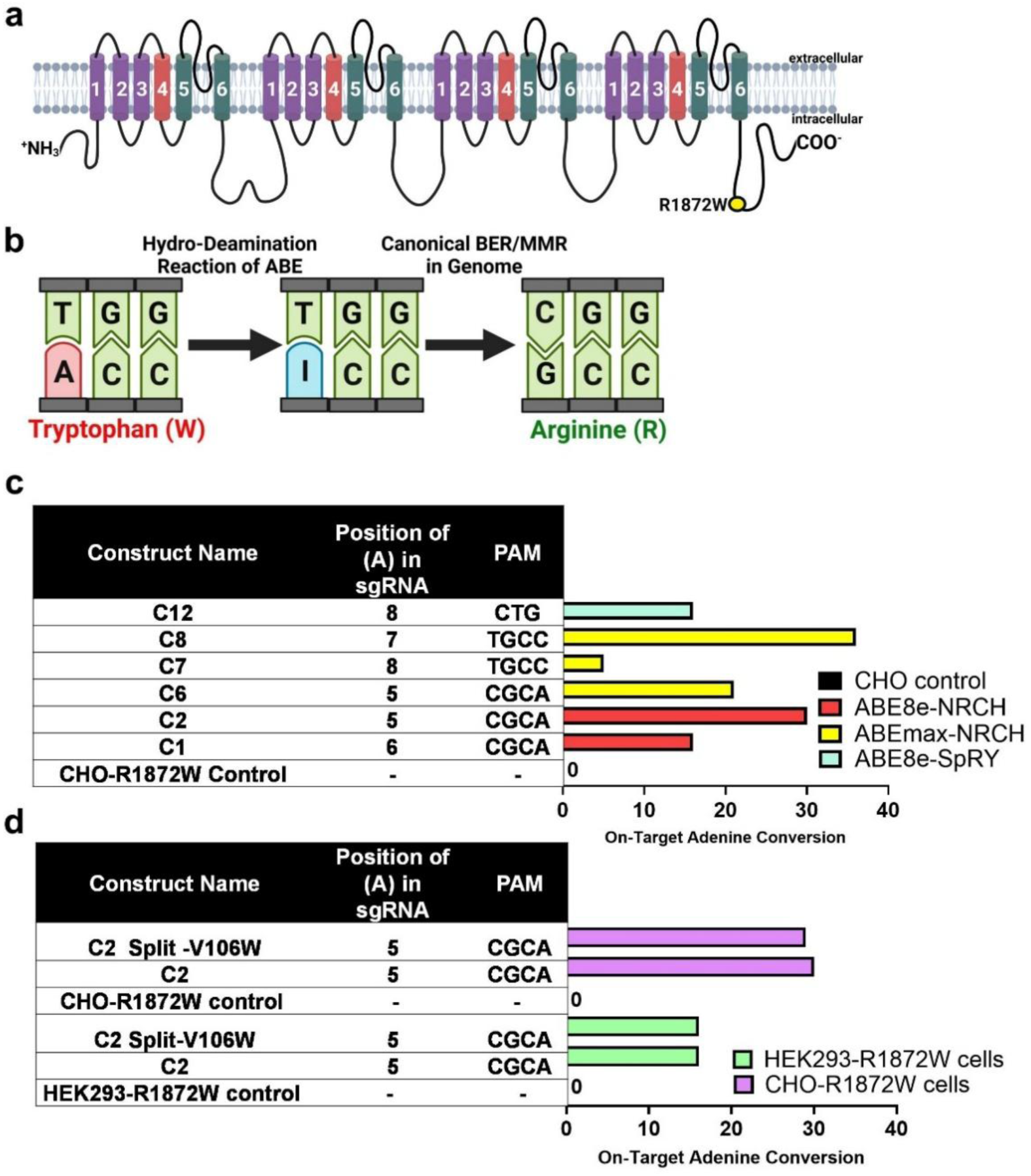
Comparative analysis of on-target ABE efficiencies in cell lines. (**a**) Location of the R1872W variant (yellow) in the C-terminal domain of the Na_v_1.6 voltage gated sodium channel. (**b**) Adenine base editing (ABE) at the R1872W locus. Target adenine (red) at the first position of the tryptophan codon is converted to inosine (blue) by the ABE and subsequently to guanine (green) via base excision and mismatch repair. Watson-Crick base pairing results in the complimentary T to C change. (**c**) Multiple constructs were screened in CHO-R1872W rodent cells. The efficacy of various base editors for on-target adenine conversion was assessed by Sanger sequencing. C2 with its paired editor ABE8e-NRCH demonstrated high efficiency targeting of the R1872W variant in the mouse genome. (**d**) Construct C2 was optimized for high on-target adenine conversion rates and minimal off-target activity in CHO-R1872W cells and HEK293-R1872W cells with the addition of a split intein structure and a V106W mutation in the TadA-8a domain. All outcomes were assessed by Sanger sequencing.

To screen for efficient base editing, sgRNA constructs were initially tested in HEK293 cells engineered to harbor a mouse-codon-optimized *SCN8A* R1872W allele (HEK293-R1872W). These cells were selected for their ease of transfection and high editing efficiencies, which were determined in our experiment by genomic DNA extraction and Sanger sequencing without the need for antibiotic selection or flow cytometry GFP sorting. Among the 16 tested constructs, 5 constructs achieved high A-to-G conversion (≥15%), while 3 constructs exhibited off-target adenine editing exceeding 2%, disqualifying them from further studies (Supp. Fig. 1). The remaining 8 constructs exhibited no significant editing of the *SCN8A* R1872W allele (Supp. Fig. 1). To mitigate the effects of HEK293 cell triploidy and account for species-specific genomic codon differences influencing guide RNA efficiency, the most effective constructs with no off-target activity were evaluated in CHO Flp-In cells. These cells, engineered to carry the same *SCN8A* R1872W allele, provide a diploid genomic context and serve as a rodent model proxy for in vivo applications (CHO-R1872W). Editing efficiencies in CHO-R1872W cells were similarly high, with 5 constructs achieving high A-to-G conversion (≥15%) (Fig. 1.c).

Construct C2 exhibited robust activity in the murine genome. We selected C2 for in vivo analyses since its paired editor, ABE8e-NRCH, demonstrated enhanced fidelity compared to prior ABEs^17^ along with compatibility to undergo mutational evolution for recognition of non-canonical PAM sequences such as NRCH^19^, where N = any nucleotide, R = A or G, and H = A, C, or T. The catalytic domain of ABE8e is a TadA deaminase that facilitates a hydro-deamination reaction, changing adenine to inosine^17^. The introduction of a V106W mutation into the TadA deaminase domain of ABE8e achieves significantly reduced DNA off-target effects^17^. The TadA-8e V106W domain is particularly advantageous in gene editing scenarios in which minimizing off-target activity is crucial, such as with the R1872W mutation, where the presence of multiple adenines in the target sequence poses a heightened risk in a sensitive pediatric population.

While the V106W mutation has been evaluated in ABE8e^17^, it has not been tested in ABE8e-NRCH. To address this, we performed site-directed mutagenesis on the TadA-8e domain of ABE8e-NRCH to introduce the V106W mutation into the base editor in a plasmid. The construct was then divided into N-terminal and C-terminal pAAV backbones, incorporating NpuN and NpuC intein proteins as previously described^25,26^ to form the C2 V106W split intein construct (Fig. 1.d). To assess whether these modifications affected construct activity, we screened them in the same engineered HEK293-R1872W and CHO-R1872W cell lines, comparing results to C2-transfected cells lacking these modifications. The C2 V106W split intein construct exhibited no significant reduction in R1872W locus editing compared to C2 in HEK293-R1872W and CHO-R1872W cell lines (Fig. 1.d).

### SCN8A-ABE treatment increases survival and ameliorates seizure activity in R1872W expressing mice

To assess the efficacy of our base editors in vivo, we used a mouse model that expresses the human derived R1872W *SCN8A* variant in a Cre-dependent manner (*Scn8a*^W/+^)^10^ (Fig. 2.a). Global expression of the R1872W variant via EIIa-Cre (*Scn8a*^W/+^-EIIa) leads to premature death at approximately P15^10^, however, peak expression of AAVs, and therefore editing activity, occurs around 3 weeks^27,28^. Due to the limited timeline for base editing in *Scn8a*^W/+^-EIIa mice, we used EMX1-Cre to generate another *SCN8A* DEE mouse model expressing the R1872W variant exclusively in forebrain excitatory neurons (*Scn8a*^W/+^-EMX1)^10^. Both mouse models typically exhibit spontaneous seizures and SUDEP, but *Scn8a*^W/+^-EMX1 mice exhibit a typical seizure onset around P20^10,29^, allowing adequate time for peak expression of our base editing AAVs and correction of the variant.

**Figure 2:**
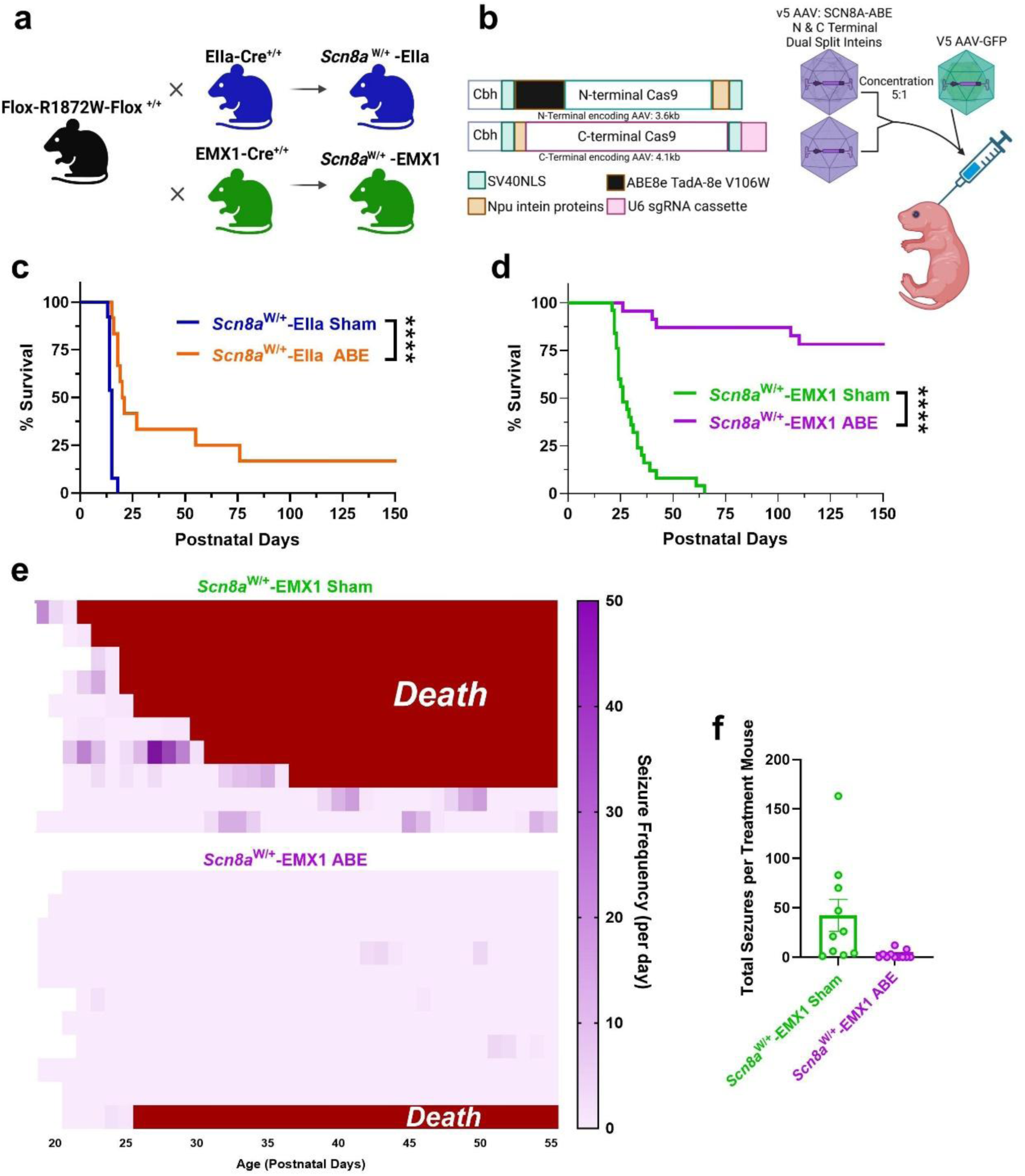
Treatment with *SCN8A*-ABE significantly increases survival and reduces seizure frequency. (**a**) Breeding strategy to generate *Scn8a*^W/+^-EIIa and *Scn8a*^W/+^-EMX1 mice. (**b)** P2 mice were injected intraventricularly (ICV) with a 5:1 ratio of AAV *SCN8A*-ABE to AAV-GFP. *Scn8a*^W/+^-EIIa and *Scn8a*^W/+^-EMX1 mice were injected with a total of 1.1 × 10^11^ viral genomes (vg) of the *SCN8A*-ABE treatment (5.5 × 10^10^ vg of each dual intein PhP.eB-ABE vectors along with 2.0 × 10^10^ vg PhP.e-GFP). Littermates of both mutant genotypes were injected only with 2.0 × 10^10^ vg of PhP.e-GFP as a ‘Sham’ viral transduction control. (**c**) Survival of *Scn8a*^W/+^-EIIa mice is significantly increased with *SCN8A*-ABE treatment (orange, n=12) compared with Sham-treated *Scn8a*^W/+^-EIIa mice (blue, n=13, *P*<0.0001, Log-rank Mantel-Cox test). (**d**) Survival of *Scn8a*^W/+^-EMX1 mice is significantly increased with *SCN8A*-ABE treatment (purple, n=24) compared with Sham-treated *Scn8a*^W/+^-EMX1 mice (green, n=25, *P*<0.0001, Log-rank Mantel-Cox test). (**e**) Seizure incidence in Sham-treated (n=10) and *SCN8A*-ABE-treated (n=11) *Scn8a*^W/+^-EMX1 mice over a period of 35 days. A red bar indicates seizure-induced death. (**f**) Sham-treated *Scn8a*^W/+^-EMX1 mice (n=10) exhibit significantly more seizures than their *SCN8A*-ABE treated counterparts (n=11); (****, *P*<0.0001, Mann-Whitney test). Data represent mean ± S.E.M.

To achieve in vivo base editing of the mutant R1872W allele, we designed a dual intein adeno-associated virus (AAV) delivery strategy, referred to as *SCN8A*-ABE, to package the split ABE8e-NRCH (V106W) base editor paired with the C2 sgRNA (C2 Split-V106W) into two AAV capsids as previously described^26^ (Fig. 2.b). The *SCN8A*-ABE treatment of dual intein vectors was paired with a third AAV directing GFP expression to visualize treated cells in later experiments. For packaging, we selected the AAV.PhP.eB capsid, an engineered variant of AAV9, due to its enhanced ability to penetrate the blood-brain barrier^30,31^. In the cortex and hippocampus specifically, previous studies show that AAV9 and PhP.eB exhibit no significant difference in neuronal tropism or in the quantity of neurons targeted when injected intracerebroventricularly (ICV) in mice^30–32^. Both serotypes have been shown to almost exclusively target neurons, and ICV injection results in efficient (>80%) transduction of neurons compared to other cells^30–32^. Given these characteristics, we utilized PhP.eB to deliver the *SCN8A*-ABE treatment along with the partnered GFP virus. As a control, we used the partnered GFP virus in the absence of *SCN8A*-ABE, referred to as Sham treatment^26^.

In both *Scn8a*^W/+^-EIIa and *Scn8a*^W/+^-EMX1 mice, we delivered the *SCN8A*-ABE or Sham treatment at P2 via ICV injection (Fig. 2.b). In Sham-treated *Scn8a*^W/+^-EIIa mice lacking the *SCN8A*-ABE, global expression of the R1872W *SCN8A* variant led to premature death at an average age of P14.5, typically following a single seizure event (Fig. 2.c). Despite the narrow timeline for efficient base editing which occurs around 3 weeks^27,28^, *SCN8A*-ABE treatment significantly increased the average survival of approximately half of all *SCN8A*-ABE treated *Scn8a*^W/+^-EIIa mice (Fig. 2.c). Two ABE-treated mice were euthanized for sequencing at P163 and P283, therefore underestimating the true survival time (Supp. Fig 2.a). In *Scn8a*^W/+^-EMX1 mice, *SCN8A*-ABE treatment significantly prolonged the survival of approximately 87% of ABE-treated mice compared to Sham-treated *Scn8a*^W/+^-EMX1 mice. Only 3 of the 24 ABE-treated *Scn8a*^W/+^-EMX1 mice succumbed to early death (>P65) (Fig. 2.d). For sequencing purposes, 16 of a total 25 *Scn8a*^W/+^-EMX1 mice treated with *SCN8A*-ABE were euthanized after P150 and another 5 after P345 (Supp. Fig 2.c), making the true survival rates for *SCN8A*-ABE-treated *Scn8a*^W/+^-EMX1 mice impossible to accurately calculate. DNA in two *SCN8A*-ABE–treated *Scn8a*^W/+^-EMX1 mice was unrecoverable.

Increased survival in *SCN8A*-ABE-treated mice was likely attributed to a decrease in seizure burden. To assess this, we evaluated seizures using concurrent video and electroencephalogram (EEG) recordings in *Scn8a*^W/+^-EMX1 mice treated either with *SCN8A*-ABE or the Sham virus (Fig. 2.e). Due to the premature death of untreated *Scn8a*^W/+^-EIIa mice, we could not record EEG signals in these mice. Spontaneous seizures were detected in all Sham-treated *Scn8a*^W/+^-EMX1 mice, and 8 of 10 mice succumbed to seizure-induced death before P40. The 2 remaining Sham-treated *Scn8a*^W/+^-EMX1 mice succumbed to death before P65. In contrast, spontaneous seizures were completely inhibited in 7 of the 11 *SCN8A*-ABE treated *Scn8a*^W/+^-EMX1 mice (Fig. 2.e, f). In another 3 mice, seizure frequency was significantly reduced (Fig. 2.e, f). We euthanized 9 of 11 mice treated with *SCN8A*-ABE at varying timepoints after P150 for genomic analysis (Supp. Fig. 2.c).

### ABE treatment demonstrates high R1872W loci targeting efficacy

We used Next-Generation targeted amplicon sequencing (NGS) to assess the extent of correction of R1872W variant alleles from mice used to generate the survival curves. We dissected hippocampal and cortical regions from 11 ABE-treated *Scn8a*^W/+^-EIIa mice and 9 of the Sham-treated *Scn8a*^W/+^-EIIa mice from the survival curve shown in Fig. 2.c. We observed a 22.2% reversion in the mutant allele (Fig. 3.a) of T/A to wildtype C/G in genomic DNA in dissected hippocampus and cortex of 6 *SCN8A*-ABE treated *Scn8a*^W/+^-EIIa mice (Fig. 3.b, Supp. Fig. 2.a, e) that survived past P21, a timepoint where all Sham-treated *Scn8a*^W/+^-EIIa mice had succumbed to death. This was detected via PCR primers spanning TALEN-silent mutations. The percentage of reads from all cells in these regions with the variant T/A at the R1872W locus of *SCN8A* was reduced from an average of 93.1% (with calculated 6.9% failure of heterozygous EIIa-Cre activation of the R1872W variant) to 70.9%. Wildtype base pairing of C/G at the R1872W locus increased with an equivalent percentage to the decrease in mutant T/A in ABE-treated *Scn8a*^W/+^-EIIa mice.

**Figure 3:**
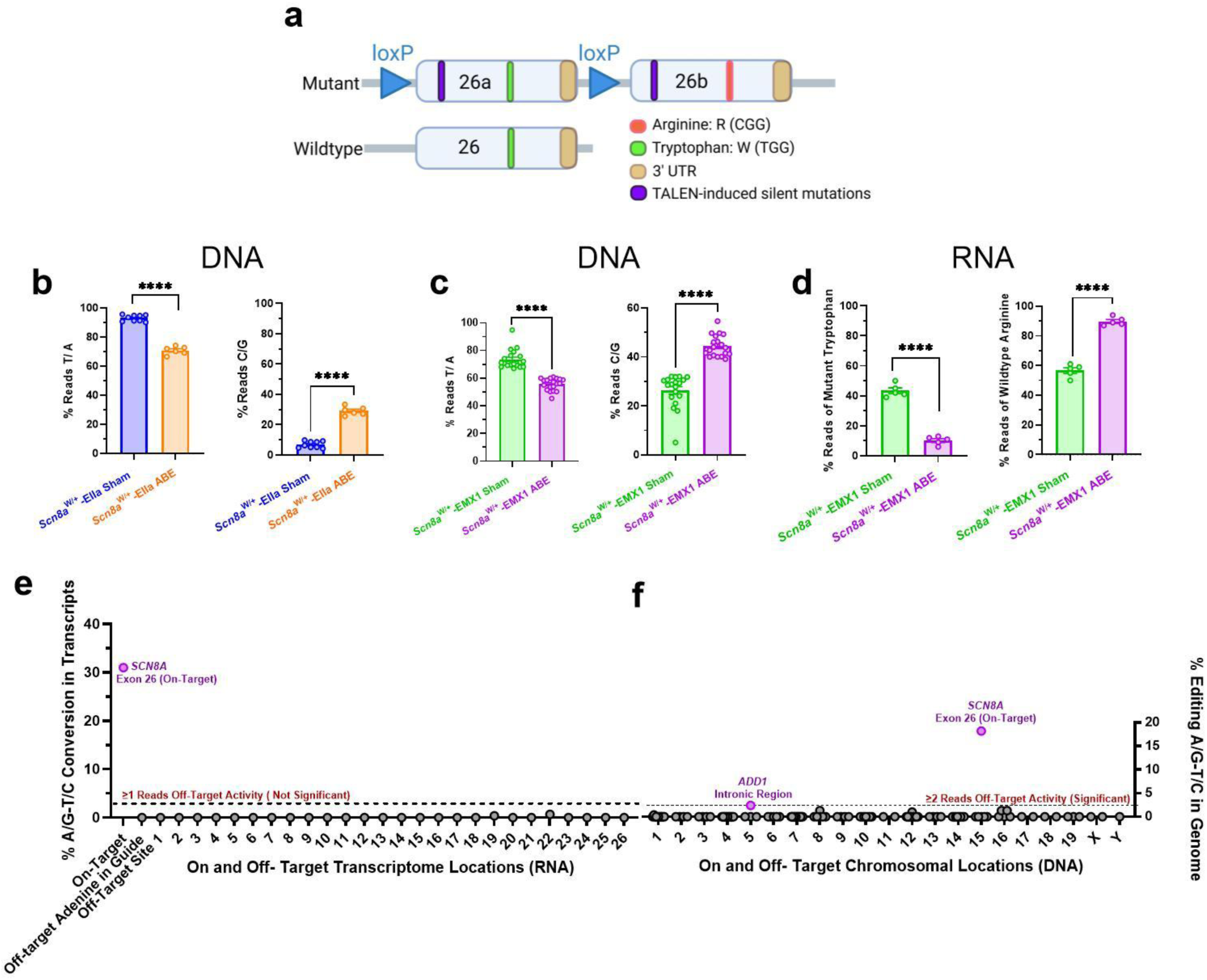
In vivo correction of the mutant T in tryptophan codon R1872W. (**a**). The mutant chromosome carries two copies of the last coding exon, designated 26a and 26b, which differ in codon 1872^10^. Exons 26a and 26b carry 8 silent base substitutions (purple) and loxP sites flanking exon 26a (triangles). DNA sequencing was carried out on fragments amplified by primers FP1 and FP2. ABE-mediated T-to-C conversion of the mutant *SCN8A*-R1872W allele in *Scn8a*^W/+^ mice restores the Wildtype allele. DNA and RNA were isolated from hippocampal and cortical dissection and homogenized for genomic processing. (**b**) Percentage of mutant T/A (left) and wildtype C/G (right) conversion in the genome of *Scn8a*^W/+^-EIIa ABE (n=6, blue) and Sham (n=9, orange) mice (****, *P*< 0.0001; Welch’s t-test). Each data point represents an individual mouse showing mutant base pair identity. (**c**) Percentage of mutant T/A (left) and wildtype C/G (right) conversion in the genome of *Scn8a*^W/+^-EMX1 ABE (n=19, purple) and Sham (n=19, green) mice (****, *P*< 0.0001; left, Mann-Whitney test; right, unpaired t-test). Each data point represents an individual mouse showing base pair identity. (**d**) Percentage of mutant tryptophan (left) to wildtype arginine (right) conversion based on transcriptome-wide RNA-seq reads in *Scn8a*^W/+^ -EMX1 ABE (n=5, purple) and Sham (n=5, green) mice (****, *P*< 0.0001; unpaired t-test). Each data point represents an individual mouse showing R1872W codon identity. (**e**) Transcriptional landscape of ABE-related off target editing in 27 of the most likely off-target loci of C2 construct. RNA-seq reads in *Scn8a*^W/+^-EMX1 ABE (purple, n=5) mice were compared to *Scn8a*^W/+^-EMX1 Sham (green, n=5) mice. Black circles represent reads with insignificant levels of editing (≤1 transcript reads). Purple circles represent reads with significant levels of editing (>1 transcript reads). Data is normalized against *Scn8a*^W/+^-EMX1 Sham mice. (**f**) Chromosomal landscape of ABE-related off target editing in 114 of the most likely off-target loci of C2 construct observed in WGS 60x Gb sequencing reads in *Scn8a*^W/+^-EMX1 ABE (n=3) mice when compared to *Scn8a*^W/+^-EMX1 Sham (n=2) mice. Black circles represent reads with insignificant levels of editing (≤2 reads). Purple circles represent reads with significant levels of editing (>2 reads). Data is normalized against *Scn8a*^W/+^-EMX1 Sham mice.

We also assessed the extent of correction of the R1872W mutation in *Scn8a*^W/+^-EMX1 mice. We dissected hippocampal and cortical regions from the 11 ABE-treated *Scn8a*^W/+^-EMX1 mice and 10 of the Sham-treated *Scn8a*^W/+^-EMX1 mice that were monitored via EEG in Fig. 2.e. Additionally, we dissected hippocampal and cortical regions from 11 of the ABE-treated *Scn8a*^W/+^-EMX1 mice and 9 of the Sham-treated *Scn8a*^W/+^-EMX1 mice from the Fig. 2.d survival curve. We observed an 18.2% reversion in the mutant allele of T/A to wildtype C/G in genomic DNA from the dissected hippocampus and cortex in a total of 19 *SCN8A*-ABE treated *Scn8a*^W/+^-EMX1 mice (Fig. 3.c, Supp. Fig. 2.c, f) that survived a timepoint where all Sham-treated mice (n=19) had succumbed to death (>P65; Fig. 2.d). The percentage of reads from all cells in these regions with the variant T/A at the R1872W locus of *SCN8A* was reduced with wildtype base pairing of C/G at the R1872W locus increased with an equivalent percentage to the decrease in mutant T/A in ABE-treated *Scn8a*^W/+^-EMX1 mice (Fig. 3.c, Supp. Fig. 2.c, f).

In both *Scn8a*^W/+^-EIIa and *Scn8a*^W/+^-EMX1 mice, *SCN8A*-ABE treatment effectively converted the tryptophan-encoding TGG sequence of the mutant allele at the R1872W locus to its wildtype arginine-encoding CGG counterpart, thereby inducing increasing levels of healthy wildtype transcript. Sequencing revealed that the editing in the hippocampus and cortex of 3 *Scn8a*^W/+^-EMX1 mice, which exhibited premature death at P26, P40, and P44, was less than 5% (Supp Fig 2. c). Thus, the observed level of editing in these mice was insufficient to support survival.

### In vivo RNA off-target analysis of SCN8A-ABE treated mice

Transcriptome-wide RNA sequencing (RNA-seq) was performed on 5 *SCN8A*-ABE treated and 5 Sham-treated *Scn8a*^W/+^-EMX1 mice. RNA-seq analysis confirmed high on-target transcript conversion with no significant bystander editing (<1%) near the target adenines (A3 and A13) following *SCN8A*-ABE treatment (Fig. 3.d,e). Sequencing of five *SCN8A*-ABE and five Sham-treated *Scn8a*^W/+^-EMX1 mice revealed no significant off-target editing (<1%) at 26 potential off-target loci. These loci, each harboring no more than five mismatches with the target sequence, were identified with CRISPOR (UCSC) ^23^, and CRISPR RGEN tools^24^, COSMID^33^ (Fig. 3.e). A total of 1,305 predicted off-target sites were considered, of which only 26 were located within exon regions expressed in the brain at the adolescent/adult mouse stage. These findings highlight the high specificity of *SCN8A*-ABE editing in vivo, further supporting its precision and translation potential.

Additionally, expression of the mutant and wild-type alleles in heterozygous *Scn8a*^W/+^-EMX1 mice confirmed equal allelic expression of both mutant and ABE-treated groups. This was accomplished by transcript detection of 2 (ATC-to-ATT) TALEN-induced silent mutations in exons 26b and 26a, which are not present in the wildtype allele ^10,34^ (Fig. 3.a, Supp. Fig. 2.g, h). This validation ensured that base editing did not alter the relative expression ratios of the heterozygous alleles, instead simply altering a single base, therefore preserving the expected transcriptional balance.

Whole genome sequencing (WGS) 60x Gb analysis in 3 *SCN8*A-ABE treated *Scn8a*^W/+^-EMX1 mice also verified high on-target DNA editing with no significant bystander editing (<2%) near the on-target adenine A3 (Fig. 3f). This was compared with 2 Sham controls. The sequencing also showed insignificant off-target editing for all but 1 of the 114 potential off-target loci with no more than three mis-match nucleotides with the target sequence predicted by CRISPOR (UCSC) ^23^, and CRISPR RGEN tools^24^, COSMID^33^. 2 of the 114 sites corresponded to protein coding genes while the other 112 sites were located within intronic/intergenic regions, including ADD1 (2.4% A to G intronic off target effects). These results again highlight the immense specificity of the *SCN8A*-ABE treatment.

### ABE treatment attenuates pathological neuronal hyperexcitability and persistent sodium current (INaP)

Increases in neuronal excitability are a hallmark of seizures in *SCN8A* DEE^11,10^, and hyperexcitability of excitatory cortical and hippocampal pyramidal cells was previously reported in the *Scn8a*^W/+^-EIIa mouse model^10^. We next determined whether *SCN8A*-ABE treatment could rescue neuronal hyperexcitability (Fig. 4). As an additional control, we examined *Scn8a*^W/+^ mice in the absence of Cre, where the R1872W variant is not activated (*Scn8a*^W/+^-control). Transduced neurons were identified by GFP expression. Fig. 4.a and b show transduction of GFP after *SCN8A*-ABE and AAV-GFP treatment in a *Scn8a*^W/+^-EMX1 mouse at P23, supporting widespread viral transduction in the brain, particularly in the hippocampus and cortex. Robust transduction was observed across cortical layers I–VI (Fig. 4.b) We examined neuronal excitability of GFP-positive pyramidal neurons in the somatosensory cortex layer IV/V of P17-25 Sham and ABE-treated *Scn8a*^W/+^-EMX1 mice. Sham-treated *Scn8a*^W/+^-EMX1 neurons were hyperexcitable when compared to *Scn8a*^W/+^-control neurons, which was attenuated in *SCN8A*-ABE treated *Scn8a*^W/+^-EMX1 neurons. *SCN8A*-ABE treatment did not fully restore firing frequencies to levels observed in *Scn8a*^W/+^-control mice at higher current injection steps (Fig. 4.c,d). Analysis of membrane and action potential (AP) properties showed a significant increase in downstroke velocity in ABE-treated *Scn8a*^W/+^-EMX1 neurons compared to Sham (Supp. Table 2). All other parameters were unchanged between ABE and Sham-treated *Scn8a*^W/+^-EMX1 neurons.

**Figure 4:**
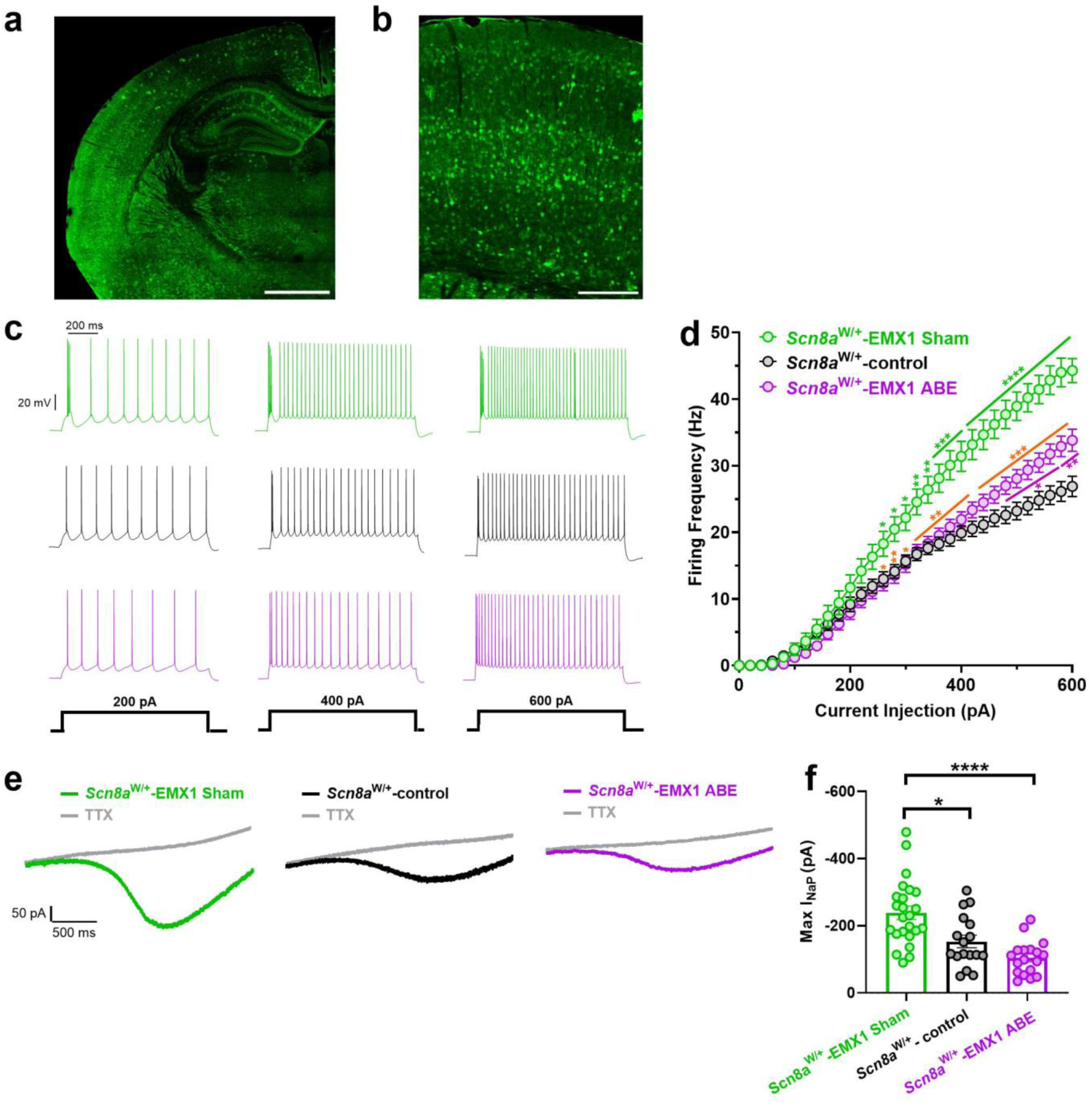
*SCN8A*-ABE attenuates neuronal hyperexcitability and reduces the pathological persistent sodium current (I_NaP_). (**a**) GFP expression in 30 µm sectioned brain tissue from *Scn8a*^W/+^-EMX1 mouse injected with the *SCN8A*-ABE and AAV-GFP viruses. Scale bar 0.5 mm. (**b**) GFP labeling of cortical layers I– VI in *Scn8a*^W/+^-EMX1 mouse injected with *SCN8A*-ABE and AAV-GFP virus. Scale bar 0.2 mm. (**c)** Example traces of action potential firing from *Scn8a*^W/+^-EMX1 Sham (green), *Scn8a*^W/+^-control (black), and *Scn8a*^W/+^-EMX1 ABE (purple) cortical layer IV/V pyramidal neurons at current injection steps of 200 pA, 400 pA, or 600 pA. (**d**) Average number of action potentials elicited relative to current injection magnitude. At current injections >240 pA, *Scn8a*^W/+^-EMX1 Sham neurons (n=25, 4 mice) exhibit increased firing frequencies when compared to *Scn8a*^W/+^-control (n=21, 4 mice). This hyperexcitability is partially attenuated in *Scn8a*^W/+^-EMX1 ABE neurons (n=34 cells, 7 mice) but *Scn8a*^W/+^-EMX1 ABE neurons exhibited significantly higher firing frequencies compared to *Scn8a*^W/+^-control at current injection steps >480 pA (*, *P*<0.05, **, *P*<0.01,***, *P*<0.001, ****, *P*<0.0001; Two-way ANOVA with Dunns’s multiple comparisons test). Green stars indicate significance between *Scn8a*^W/+^-EMX1 Sham and *Scn8a*^W/+^-control groups and purple stars indicate significance between *Scn8a*^W/+^-EMX1 ABE and *Scn8a*^W/+^-control groups. Orange stars indicate significance between *Scn8a*^W/+^-EMX1 ABE and *Scn8a*^W/+^-EMX1 Sham groups. (**e**) Example traces of steady state I_NaP_ evoked by slow voltage ramps in pyramidal neurons from *Scn8a*^W/+^-EMX1 Sham (green), *Scn8a*^W/+^-control (black), and *Scn8a*^W/+^-EMX1 ABE mice (purple). Traces in gray show slow voltage ramp in the presence of 500 nM TTX. (**f**) Elevated maximum I_NaP_ in *Scn8a*^W/+^-EMX1 Sham neurons (n=24, 7 mice) is rescued by ABE treatment in *Scn8a*^W/+^-EMX1 neurons (n=17, 8 mice) to amplitudes recorded in *Scn8a*^W/+^-control neurons (n=17, 6 mice), (*, *P*<0.05, ****, *P*<0.0001; Kruskal-Wallis with Dunn’s multiple comparisons). No significant difference observed between *Scn8a*^W/+^-EMX1 ABE and *Scn8a*^W/+^-control groups. Data represents mean ± S.E.M.

We also observed a heightened neuronal excitability in P13-17 *Scn8a*^W/+^-EIIa Sham mice when compared to *Scn8a*^W/+^-control, which was completely rescued by *SCN8A* ABE-treatment (Supp Fig. 3.a, b). Analysis of membrane and action potential properties revealed a decrease in rheobase and downstroke velocity and an increase in AP width in *Scn8a*^W/+^-EIIa Sham mice when compared to *Scn8a*^W/+^-EIIa ABE mice. We also observed a decrease in AP threshold in both Sham and ABE-treated *Scn8a*^W/+^-EIIa mice when compared to *Scn8a*^W/+^-control. (Supp. Table 3).

The most prominent biophysical feature of *SCN8A* GOF variants is an elevated persistent sodium current (I_NaP_)^3,11,35^. Elevated I_NaP_ is a major contributor to neuronal hyperexcitability in epilepsy^35^. Thus, we sought to determine whether *SCN8A*-ABE treatment eliminated the pathological heightened I_NaP_. We used slow voltage ramps in pyramidal neurons from layer IV/V of the somatosensory cortex to measure I_NaP_ in Sham and *SCN8A*-ABE-treated *Scn8a*^W/+^-EMX1 mice and compared them to *Scn8a*^W/+^-control mice (P17-P25). ABE treatment significantly decreased the amplitude of I_NaP_ compared with Sham-treated *Scn8a*^W/+^-EMX1 mice to levels recorded in *Scn8a*^W/+^-control neurons (Fig. 4.e, f). Half maximal activation voltages (V_1/2_) were not different between the three groups (Supp. Table 2).

### ABE treatment improves behavioral comorbidities in R1872W SCN8A DEE mice

In addition to seizures, *SCN8A* DEE patients suffer from severe comorbidities including movement disorders and cognitive impairment^5^. To examine locomotor activity and anxiety-like behavior, we evaluated both Sham and *Scn8a*-ABE-treated *Scn8a*^W/+^-EMX1 mice along with *Scn8a*^W/+^-control mice in the open field test. We used total distance travelled to assess locomotor activity and thigmotaxis as a measure of anxiety-like behavior^36^. Mice were tested between 4 and 8 weeks of age, a time point after the onset of seizures in *Scn8a*^W/+^-EMX1 mice^10^. We generated quantifications of locomotor activity to analyze behavior in all groups (Fig. 5). *Scn8a*^W/+^-EMX1 Sham mice had reduced locomotor activity and spent significantly less time in the open areas of the field compared to *Scn8a*^W/+^-control mice, indicating a potential motor impairment and increased anxiety (Fig. 5.a-c). Treatment with *SCN8A*-ABE significantly improved both comorbidities, with locomotor activity restored to levels observed in *Scn8a*^W/+^-control mice (Fig. 5b). Thigmotaxis was significantly reduced in *SCN8A*-ABE-treated *Scn8a*^W/+^-EMX1 mice, but did not reach *Scn8a*^W/+^-control levels (Fig. 5.a,c). These findings collectively suggest that *Scn8a*^W/+^-EMX1 mice have underlying motor impairments and anxiety that can be ameliorated by early *SCN8A*-ABE treatment.

**Figure 5:**
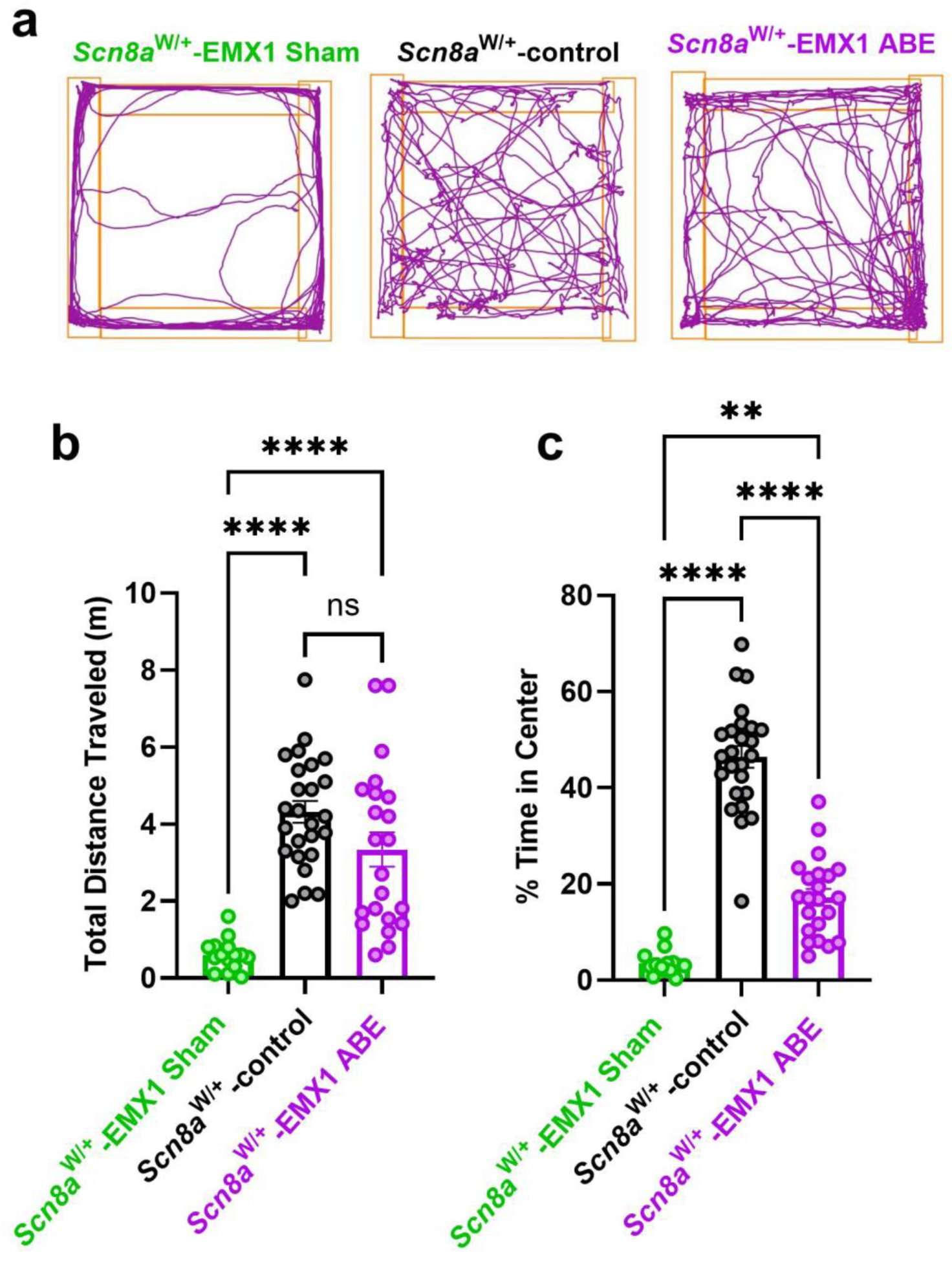
Anxiety and movement disorder phenotypes are reduced by *SCN8A*-ABE treatment. (**a**) Representative movement tracks in open field for *Scn8a*^W/+^-control (n=25), *Scn8a*^W/+^-EMX1 Sham (n=15), and *Scn8a*^W/+^-EMX1 ABE mice (n=22). (**b**) In the open field test, *Scn8a*^W/+^-EMX1 Sham mice (n=15) show significantly decreased distance travelled compared to *Scn8a*^W/+^-control (n=25). Treatment with *SCN8A*-ABE in *Scn8a*^W/+^-EMX1 mice (n=22) rescues this phenotype to levels seen in *Scn8a*^W/+^-control mice (****, *P*<0.0001, Brown-Forsythe ANOVA with Dunnett’s multiple comparisons test). (**c**) *Scn8a*^W/+^-EMX1 Sham mice (n=15) show significantly reduced time in the center in the open field test compared to *Scn8a*^W/+^-controls (n=25). *Scn8a*^W/+^-EMX1 ABE treatment attenuated this phenotype (n=22) but did not reach *Scn8a*^W/+^-control levels (**, *P*<0.01, ****, *P*<0.0001, Kruskal-Wallis test with Dunn’s multiple comparisons test). Data represents ± S.E.M.

## Discussion

Treatments for severe genetic epilepsy disorders are limited, and current therapeutic approaches do not target the underlying mechanisms. Here, we provide compelling evidence that base editors can serve as an effective and targeted therapeutic approach for one *SCN8A* DEE variant, R1872W, validating the potential for targeted treatment of other *SCN8A*-related DEE variants.

Our results show that ABE treatment successfully reduced seizure burden of mice carrying the *SCN8A* R1872W variant, which increased survival and improved associated behavioral comorbidities. Neuronal hyperexcitability, a hallmark of seizure activity, and elevation of persistent sodium current (I_NaP_) associated with some *SCN8A* GOF variants^11,35^ were also attenuated in ABE-treated mice. These findings support a reversal of the pathogenic GOF physiology of the R1872W variant via *SCN8A*-ABE treatment. This substantial phenotypic rescue was achieved by correction of approximately 30% of the mutant *SCN8A* transcripts, which is primarily expressed by neurons^2,37^. RNA sequencing revealed no significant off-target effects, supporting high specificity of editing of the R1872W base. The use of an optimized guide RNA along with a highly specific evolved ABE8e base editor minimized off-target effects while preserving high editing efficiency. These findings underscore the transformative potential of precise gene-editing interventions in genetic epilepsy disorders where current pharmacological treatments remain ineffective.

Our study is the first to use base editing in genetic forms of epilepsy, but other gene therapies have been explored previously as a treatment for *SCN8A* DEE, Dravet Syndrome, and other severe genetic epilepsy syndromes. Antisense oligonucleotides (ASOs) ameliorate many aspects of the DEE phenotype in mice^38–42^, and approximately 25% reduction of *SCN8A* transcripts has been shown to rescue the premature death phenotype of *SCN8A* DEE^39^. However, ASOs require continuous administration due to turnover within a few weeks to months of injection^43^. Furthermore, the turnover rate of Na_v_1.6 transcripts, which is not completely understood^1^, could also impact the success of ASOs in *SCN8A* DEE^44^. In contrast, base editing can offer permanent treatment to revert pathogenicity by directly targeting the genetic cause of the *SCN8A* DEE phenotype. In this study, we achieved a 30% permanent reversion of mutant to wild-type transcripts, indicating that our editing efficiency is more than sufficient for phenotypic rescue.

Allele-specific editing using CRISPR/Cas9 has been explored as a therapeutic strategy for *SCN8A* DEE in mice and demonstrated the feasibility of specifically inactivating the pathogenic *SCN8A* variant^45^. By using silent mutations to create sequence differences between mutant and wild-type alleles, selective targeting of the mutant allele was achieved. However, while this strategy was effective in mice engineered to ubiquitously express Cas9, translating it to human patients is more complex due to the absence of natural sequence variation and challenges in Cas9 delivery. Nonetheless, this study provides a critical foundation, demonstrating that CRISPR-based modifications of sodium channel genes can drive functional changes, guiding future therapeutic endeavors.

The most striking effect of *SCN8A*-ABE treatment was the profound inhibition of seizure activity observed in the majority of ABE-treated *Scn8a*^W/+^-EMX1 mice and a significant reduction in seizure frequency in the remaining mice. This directly translated to increased survival in *Scn8a*^W/+^-EMX1 mice and likely the same for the ABE-treated *Scn8a*^W/+^-EIIa mice. In addition to seizures, *SCN8A* DEE patients experience severe comorbidities, including motor impairments and anxiety-like behaviors^46,47^. Our behavioral analyses revealed that *SCN8A*-ABE treatment significantly improved locomotor activity and reduced anxiety-like behaviors in *Scn8a*^W/+^-EMX1 mice. This suggests that restoring early Na_v_1.6 function may have therapeutic benefits beyond seizure suppression, addressing other debilitating aspects of the disorder.

The physiological consequences of *SCN8A* GOF variants include an elevated I_NaP_ due to disrupted channel inactivation and subsequent neuronal hyperexcitability ^3,10,11^. Consequentially, many pharmacological treatments for *SCN8A* DEE target I_NaP_ directly ^48,49^. *SCN8A-*ABE treatment ameliorated the pathological increase in I_NaP_ in cortical neurons to non-pathological levels recorded in control mice, which likely led to the rescue of neuronal excitability in ABE-treated cortical neurons. Expression of the *SCN8A* R1872W variant selectively in cortical neurons is sufficient for seizure onset and seizure-induced death^10^. This suggests that focused targeting of key areas of the brain, namely the hippocampus and cortex, could be sufficient for seizure suppression.

We observed higher on-target RNA levels of reversion (approximately 30%) of the mutant expression than observable in DNA (approximately 20%) following the delivery of the AAV vector PhP.eB. This observation simply reflects the preferential transfection of neurons by PhP.eB and demonstrates the phenomenon of the *SCN8A* gene being preferentially expressed in neurons in the CNS^2,37^.

A major concern in gene-editing applications is the potential for off-target modifications that could result in unintended consequences. RNA transcriptome-wide analysis revealed high specificity of *SCN8A*-ABE, with no significant DNA or RNA off-target mutations detected. This is likely due to the high specificity of the guide RNA as well as the use of ABE8e[TadAV106W]-NRCH. ABE8e itself demonstrates a minor degree of off-target editing which was ameliorated with the addition of variant V106W in the TadA deaminase domain of ABE8e^17^. We elected to induce the V106W variant via site directed mutagenesis (SDM) into ABE8e-NRCH to achieve off-target reductions. Studies have demonstrated that adenine base editors have the ability to cause off-target effects within the genome^50^. Unintended modifications at non-target sites pose significant challenges for clinical applications of all genome editors. These unintended mutations can lead to disruption of essential genes or the activation of oncogenes, potentially resulting in severe side effects like tumor development^51^. To mitigate these risks, it is crucial to develop strategies like the V106W variant incorporation and highly specific guide RNAs that enhance the precision of base editors.

While this study provides strong evidence for the therapeutic potential of base editing in *SCN8A* DEE, several challenges remain. The observed editing efficiency, while sufficient for phenotypic rescue, remains below the theoretical maximum. Future strategies to enhance editing, such as optimizing AAV dosing with dual guide RNA exposure^26^ or employing novel viral capsids or alternative delivery methods, such as lipid nanoparticles with improved CNS-wide targeting, may be required for full symptomatic relief in these patients. Since *SCN8A* DEE patients typically exhibit seizure onset at approximately 4 to 6 months of age and can be identified shortly after that time^52^, a major consideration for the applicability of base editing in these patients is the time of delivery. In our study, both *Scn8a*^W/+^-EIIa and *Scn8a*^W/+^-EMX1 mice were treated at postnatal day 2 before the onset of spontaneous seizures. In human patients, ABE treatment would be initiated after initial seizure onset. This is true for other genetic epilepsy syndromes as well^38,39,42,45^. Administration of an *SCN8A* ASO or shRNA after seizure onset has been shown to be protective against SUDEP in two models of *SCN8A* DEE^41^. While future studies are needed to assess the efficacy of base editing in *SCN8A* DEE after seizure onset, it is likely that a “therapeutic window” can be identified to allow AAV expression and turnover of the pathologic protein.

Patients with variants at residue R1872 constitute approximately 10-15% of reported cases of *SCN8A* DEE^4,7^. Many other recurrent GOF *SCN8A* variants such as R1617Q lead to a DEE phenotype^53^, and these could be targeted by base editors for precise therapeutic interventions that address the underlying genetic cause of the disease. Base editing technology may also be utilized to target severe single-nucleotide variants responsible for other genetic epilepsy syndromes. As gene-editing technologies continue to advance, base editing approaches such as *SCN8A*-ABE pave the way for future clinical applications of gene therapies in genetic forms of epilepsy.

## Methods

### Cell Culture

HEK293 cells stably expressing the R1872W variant were generated by the UVA Genome Engineering Shared Resource (GESR) Core. A 34-base-long double-stranded FRT sequence was introduced into the AVS1 AAV integration locus on chromosome 19 of HEK293 cells (ATCC) with CRISPR-Cas9. For stable integration of plasmids, these stably expressing FRT-HEK293 cells were cotransfected with two plasmids at an equimolar ratio as previously descibed^54^: the first being pCAG-Flp recombinase (Addgene Plasmid #13787) and the second a plasmid containing FLP sites (inserted by Genscript Inc.) flanking the *SCN8A* coding sequence with the R1872W variant^7,55^. The latter plasmid, prior to transfection, was subjected to SDM by Genscript Inc. to include the R1872W variant as well as three silent mutations^4,7,34^ that were introduced to allow for polymerase chain reaction (PCR) amplification, accomplished by primer pair FP1 and RP1, of the mutant allele only. This approach streamlined workflow and improved accuracy of on-target analysis. Consequently, the transfection mentioned above allowed for the insertion of the mouse-codon-optimized synthetic *SCN8A* sequence into the AVS1 locus in one chromosome of the HEK293 chr.19 triploid group via FLP-Recombinase/FRT directed insertion, partnered with a geneticin (G418) resistance cassette (HEK293T-R1872W). The latter enabled antibiotic selection post-integration, thereby establishing a stabilized R1872W cell line conducive to the swift screening of sgRNA sequences. A CHO cell line (Invitrogen; Flp-In™-CHO Cell Line) containing the R1872W variant was created via the same transfection protocol (CHO-R1872W). Culture of CHO-R1872W and HEK293T-R1872W cells was performed according to previously published protocols^17^. Cells were maintained in Dulbecco’s Modified Eagle medium (DMEM; Life Technologies) supplemented with 10% fetal bovine serum (Gibco), 0.1 mM nonessential amino acids (NEAA, Life Technologies), Glutamax (GM, Life Technologies).

### Cloning and Viral Packaging

The sgRNA expression plasmid pU6-pegRNA-GG-acceptor (Addgene Plasmid #132777) was subjected to mRFP1 removal and replacement of a GFP cassette partnered with a self-cleaving P2A peptide by Genscript Inc. The GFP-P2A insertion enables the guide RNA to be transcribed independently of GFP. This allowed for rapid assessment of transfection efficiency mediated by the guide RNA plasmid. Subsequent individual sgRNAs were then cloned into the sgRNA expression plasmid (sgRNA-P2A-GFP) by Genscript Inc. Constructs were transformed into Top10 chemically competent *Escherichia coli* (Thermo Fisher) grown on LB agar plates, and liquid cultures were grown in LB broth overnight at 37°C with 100 μg/ml ampicillin. Individual colonies were validated by Sanger sequencing by Genscript Inc. Verified plasmids were prepared by maxiprep (Qiagen).

### Base Editor Construct Characterization

For characterization of base editing constructs, mutation-integrated cells were seeded 16–18h before transfection. Both cell types (HEK293T-R1872W and CHO-R1872W) were subjected to transfection at approximately 70% confluency with a plasmid encoding the base editor and a separate plasmid harboring the sgRNA-P2A-GFP construct. Transfection of base editor and guide construct occurred at a 3:1 mass ratio using Lipofectamine 3000 (ThermoFisher Scientific) as previously described^17^ and in accordance with the manufacturer’s protocols, but scaled up into a 6-well plate (polystyrene, Corning). This scale-up was done to evaluate high cell density conditions in either cell line. Cells did not undergo antibiotic selection or GFP cell sorting after base editor exposure during transfection and were cultured for 72 hours before harvesting. Culture of HEK293T-R1872W and CHO-R1872W cells was performed in Dulbecco’s Modified Eagle medium (DMEM; Life Technologies) supplemented with 10% fetal bovine serum (Gibco), 0.1 mM nonessential amino acids (NEAA, Life Technologies), Glutamax (GM, Life Technologies. We collected gDNA from cells 72 hours after transfection and Genomic DNA was isolated using PurelinkDNA mini Kit (Invitrogen). PCR was performed at the following conditions:

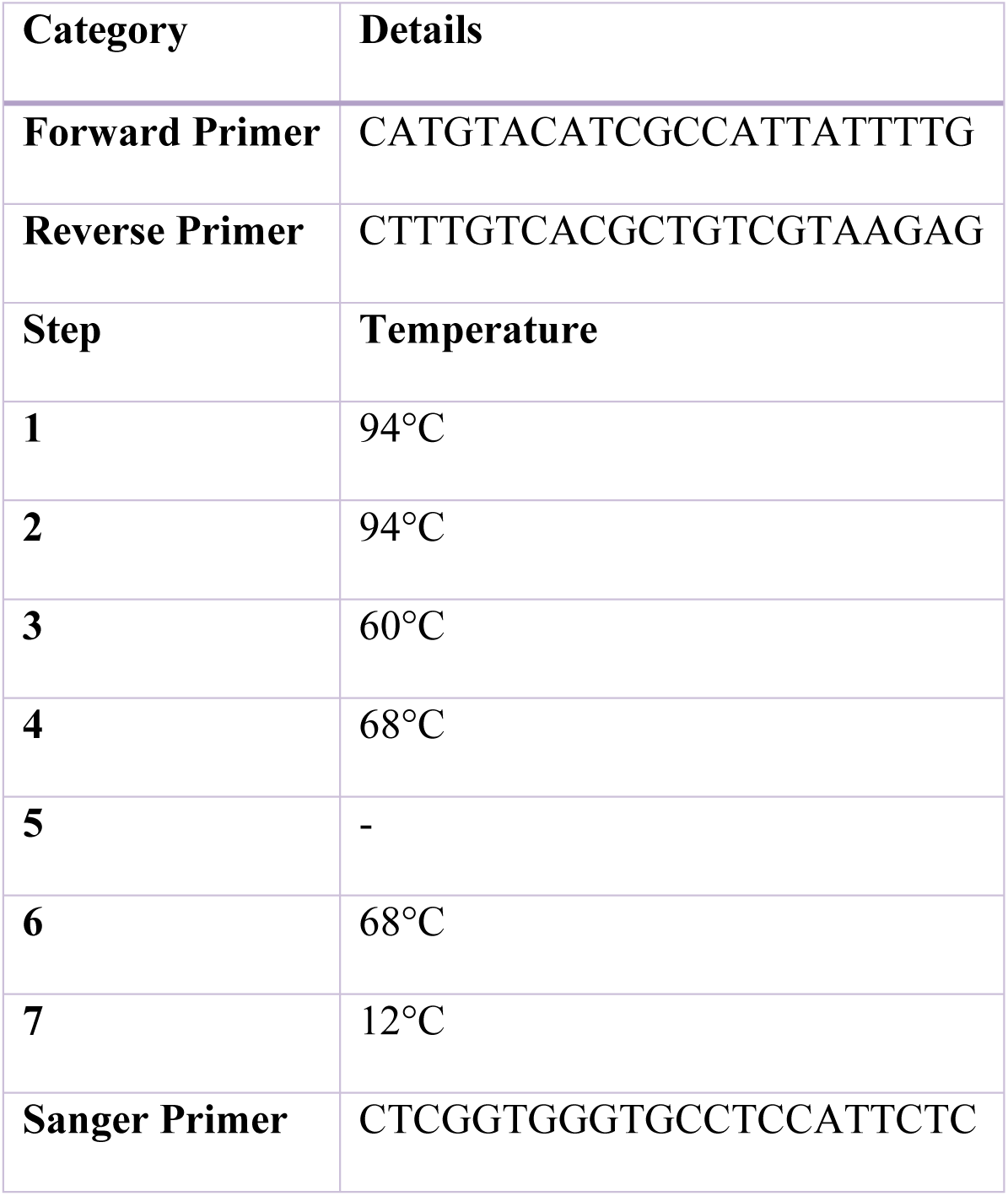

PureLink PCR Purification Kit was utilized following completion of PCR. Sanger sequencing of the purified product of edited *SCN8A* in HEK293T-R1872W and CHO-R1872W cells was performed at Eurofins Genomics and analysis of files were conducted with EditR1.0.10 as previously described^56^ and verified on SnapGene v.10.0.3.

### Mouse Husbandry and Genotyping

All experiments using live animals were approved by the University of Virginia Animal Care and Use Committees. *Scn8a*^W/+^ mice were generated as previously described^4,7^ and maintained through crosses with C57BL/6J mice (Jax, #000664) to keep all experimental mice on a C57BL/6J genetic background.^7,10^ Cell type-specific expression of R1872W was achieved using males heterozygous for the R1872W allele and C57BL/6J females homozygous for either EIIa-Cre (Jax, #017320) or EMX1-Cre (Jax 005628) to generate mutant mice (*Scn8a*^W/+^-EIIa and *Scn8a*^W/+^-EMX1).^10^ Homozygous Cre females were used for breeding to ensure minimal germline recombination due to Cre, as shown previously ^57,58^. For all experiments we used *Scn8a*^W/+^ mice as controls that contained the same allele encoding the *SCN8A* variant but lacked Cre activation and therefore only expressed wildtype Na_v_1.6 from both alleles^4^.

### Viral Vectors and Intracerebroventricular injections

AAV vector backbone plasmids were cloned by Genscript Inc. to insert the sgRNA sequence and C-terminal base editor half of ABE8e-NRCH (V106W) into v5 Cbh-AAV-ABE-NpuC+U6-sgRNA (Addgene 137177), and the N-terminal base editor half v5 Cbh-AAV-ABE-NpuN (Addgene 137178)^26^. Viral capsid preparations, packaging, and purification of PhP.eB -AAV packaged base editor plasmids were performed by Salk Institute’s Gene Transfer, Targeting and Therapeutics Viral Vector Core. High-titer qualified AAV was stored at −80°C until use. Neonatal ICV injections were performed as previously described^10^. *Scn8a*^W/+^-EIIa and *Scn8a*^W/+^-EMX1 mice were given a single unilateral ICV (2 μL volume) of either 2.0 × 10^10^ viral genomes (vg) PhP.eB-CMV-GFP in PBS (Sham) or 1.1 × 10^11^ vg of the *SCN8A*-ABE treatment in PBS (5.5 × 10^10^ vg of each dual intein PhP.eB-ABE vectors) along with 2.0 × 10^10^ vg PhP.e-GFP at P1 using a 33-gauge needle attached to a 5 μL microvolume syringe as previously described^59^. This dosage aligns with high therapeutic doses used for P0 ICV AAV administration of base editors for rescuing Hutchinson-Gilford Progeria syndrome and Niemann-Pick disease in mice^26,60^.

### Whole Brain Immunofluorescence Imaging

Brain tissue for immunohistochemistry was processed as follows: Mice were anesthetized and transcardially perfused with 10 ml ice-cold Dulbecco’s PBS (DPBS, Gibco, 14200-075) followed by 10 ml ice-cold 4% PFA. Brains were fixed in 4% paraformaldehyde (PFA) and 30 μm coronal brain sections were obtained using a cryostat. Floating sections were fixed and stored in DPBS. Sections were permeabilized with 0.1% Triton X-100 in DPBS for 30 minutes and blocked with 2% bovine serum albumin (BSA) for 2 hours. Sections were incubated with the primary antibody, rabbit anti-GFP (Abcam, ab290), diluted in DPBS containing 2% BSA at a concentration of 1:1000. Secondary antibodies were diluted 1:500 in DPBS containing 2% BSA. Sections were stained free-floating in primary antibody on a shaker at 4°C overnight and with secondary antibody for 2h at room temperature the following day. Sections were mounted using Prolong Gold antifade reagent with DAPI (Invitrogen, P36935) on a microscope slide and covered with a cover glass. Images were gathered using Zeiss Zen software on a LSM700 or LSM880. Whole brain images were stitched following acquisition with a 20x objective (Plan-Apochromat 20 x/0.8) and z-stack images were gained by use of a 63x objective (Plan-Apochromat 63 x/1.4).

| Category | Item | Catalog Number |
| --- | --- | --- |
| Primary Antibodies | Abcam Rb pAb to GFP | ab290 |
| Secondary Antibodies<br>(Alexa Fluor) | 488 goat anti-mouse IgG (H+L) | A11029 |
|  | 555 goat anti-rabbit IgG (H+L) | A21429 |
|  | 594 goat anti-rabbit IgG (H+L) | A11037 |

### Brain Slice Preparation

Preparation of acute brain slices for patch-clamp electrophysiology experiments was modified sparingly from standard protocols previously described^11^. Mice were anesthetized with isoflurane and decapitated. The brains were rapidly removed and kept in chilled slicing solution (1°C) containing (in mM): 93 N-Methyl-D-glutamine (NMDG), 2.5 KCl, 1.25 NaH_2_PO_4_, 20 HEPES, 5 L-ascorbic acid (sodium salt), 2 thiourea, 3 sodium pyruvate, 0.5 CaCl_2_, 10 MgSO_4_, 25 D-glucose, 12 N-acetyl-L-cysteine, 30 NaHCO_3_; pH adjusted to 7.4 using HCl (osmolarity 310 mOsm). Slices were continuously oxygenated with 95% O_2_ and 5% CO_2_ throughout the preparation. 300 µm coronal brain sections were prepared using a Leica Microsystems VT1200 vibratome. Slices were collected and placed in warmed (37°C) ACSF containing (in mM): 125 NaCl, 2.5 KCl, 1.25 NaH_2_PO_4_, 2 CaCl_2_, 1 MgCl_2_, 0.5 L-ascorbic acid, 10 glucose, 25 NaHCO_3_, and 2 Na-pyruvate. Slices were incubated for approximately 30 min and then kept at room temperature for up to 6 h.

### Intrinsic Excitability Recordings

Brain slices were placed in a chamber superfused (∼2 ml/min) with continuously oxygenated recording solution warmed to 32 ± 1°C. Whole-cell recordings were performed using a Multiclamp 700B amplifier with signals digitized by a Digidata 1322A digitizer. Currents were amplified, lowpass filtered at 2 kHz, and sampled at 100 kHz. Borosilicate electrodes were fabricated using a Brown-Flaming puller (model P1000, Sutter Instruments) to have pipette resistances between 3 and 5 mΩ. Current-clamp recordings of neuronal excitability were collected in ACSF solution. The internal solution contained the following (in mM): 120 K-gluconate, 10 NaCl, 2 MgCl_2_, 0.5 K_2_EGTA, 10 HEPES, 4 Na_2_ATP, 0.3 NaGTP, pH 7.2 (osmolarity 290 mOsm). Intrinsic excitability was assessed using methods adapted from those previously described^11^. Resting membrane potential was manually recorded from the neuron at rest. The first AP at rheobase, defined as the maximum amount of depolarizing current that could be injected into neurons before eliciting an AP, was used to calculate passive membrane and AP properties, including threshold, upstroke and downstroke velocity, which are the maximum and minimum slopes on the AP, respectively; amplitude, which was defined as the voltage range between AP peak and threshold; APD_50_, which is the duration of the AP at the midpoint between threshold and peak; and input resistance, which was calculated using a −20 pA pulse in current-clamp recordings. AP frequency–current relationships were determined using 1s current injections from −140 to 600 pA. All patch-clamp electrophysiology data were analyzed using custom MATLAB R24.a scripts and/or ClampFit 11.2.

### Persistent Sodium Current (INaP) Recordings

The recording solution has been previously described^11^ and contained (in mM): 100 NaCl, 40 TEACl, 10 HEPES, 3.5 KCl, 2 CaCl_2_, 2 MgCl_2_, 0.2 CdCl_2_, 4 4-aminopyridine (4-AP), 25 D-glucose. The internal solution for sodium channel recordings contained the following (in mM): 140 CsF, 2 MgCl_2_, 1 EGTA, 10 HEPES, 4 Na_2_ATP, and 0.3 NaGTP with the pH adjusted to 7.3 and osmolality to 300 mOsm. Steady-state persistent sodium currents (I_NaP_) were elicited using a voltage ramp (20 mV/s) from −80 to −20 mV. After collecting recordings at baseline, protocols were repeated in the presence of 500 nM tetrodotoxin (TTX; Alomone Labs) to completely isolate I_NaP_. TTX-subtracted traces were analyzed by subtracting the current recorded in the presence of TTX from the current recorded in its absence. The half-maximal voltage for activation was calculated as previously described^61^.

### Electroencephalogram (EEG) Recordings

Custom electroencephalogram (EEG) headsets (Digikey) were implanted in P18 *Scn8a*^W/+^-EMX1 mice using standard surgical techniques as previously described.^62^ Anesthesia was induced with 5% and maintained with 0.5%-3% isoflurane. Adequacy of anesthesia was assessed by lack of toe-pinch reflex. A midline skin incision was made over the skull and connective tissue was removed. Burr holes were made at the lateral/anterior end of the left and right parietal bones to place EEG leads, and at the interparietal bone for ground electrodes. EEG leads were placed bilaterally in the somatosensory cortex and unilaterally placed in the occipital lobe. A headset was attached to the skull with dental acrylic (Jet Acrylic; Lang Dental). Mice received postoperative analgesia with ketoprofen (5 mg/kg, i.p.) and 0.9% saline (0.5 mL i.p.) and were allowed to recover a minimum of 2-4d before seizure-monitoring experiments. Mice were then individually housed in custom-fabricated chambers and monitored for the duration of the experiment. The headsets were attached to a custom low-torque swivel cable, allowing mice to move freely in the chamber. EEG signals were amplified at 2000× and bandpass filtered between 0.3 and 100 Hz, with an analog amplifier (Neurodata model 12, Grass Instruments). Biosignals were digitized with a Powerlab 16/35 and recorded using LabChart 7 software at 1 kS/s. Video acquisition was performed by multiplexing four miniature night vision-enabled cameras and then digitizing the video feed with a Dazzle Video Capture Device and recording at 30 fps with LabChart 7 software in tandem with Biosignals.

### Next Generation Amplicon Genomic DNA Sequencing

Sequencing library preparation was performed according to previously published protocols^22^. We isolated genomic DNA (gDNA) with the Invitrogen™PureLink™ Genomic DNA Mini Kit and used approximately 25mg of tissue was used for individual locus editing experiments. DNA from hippocampal and cortical microdissection and homogenized for genomic processing. Sequencing libraries were amplified in three steps, first to amplify the mutated locus of interest to confirm editing of the mutant allele without healthy read noise from the Wildtype allele of the heterozygote mice (FP1/RP1). This was accomplished by detection of TALEN-induced silent mutations in exons 26b and 26a, which are not present in the wild-type allele^34^. The second primer set functioned to shorten the length of the amplicon to 150bp (FP2/RP2); and third to add full-length Illumina sequencing adapters:

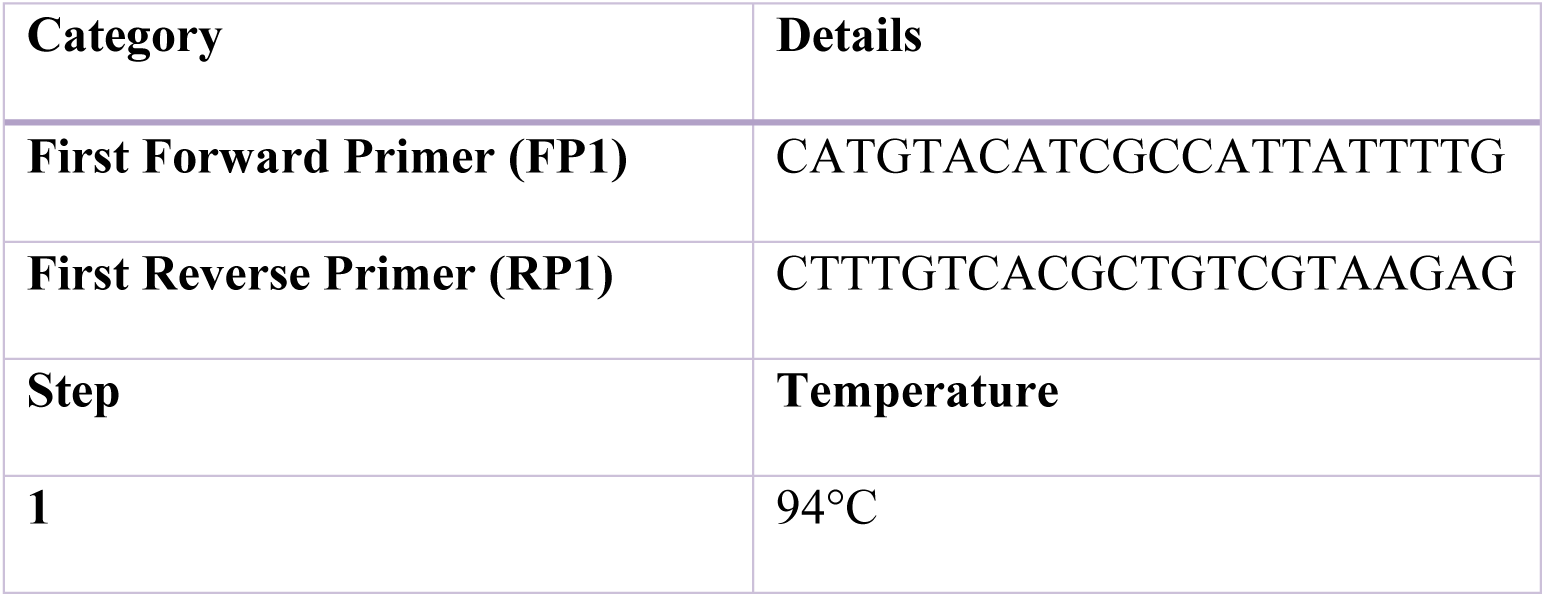
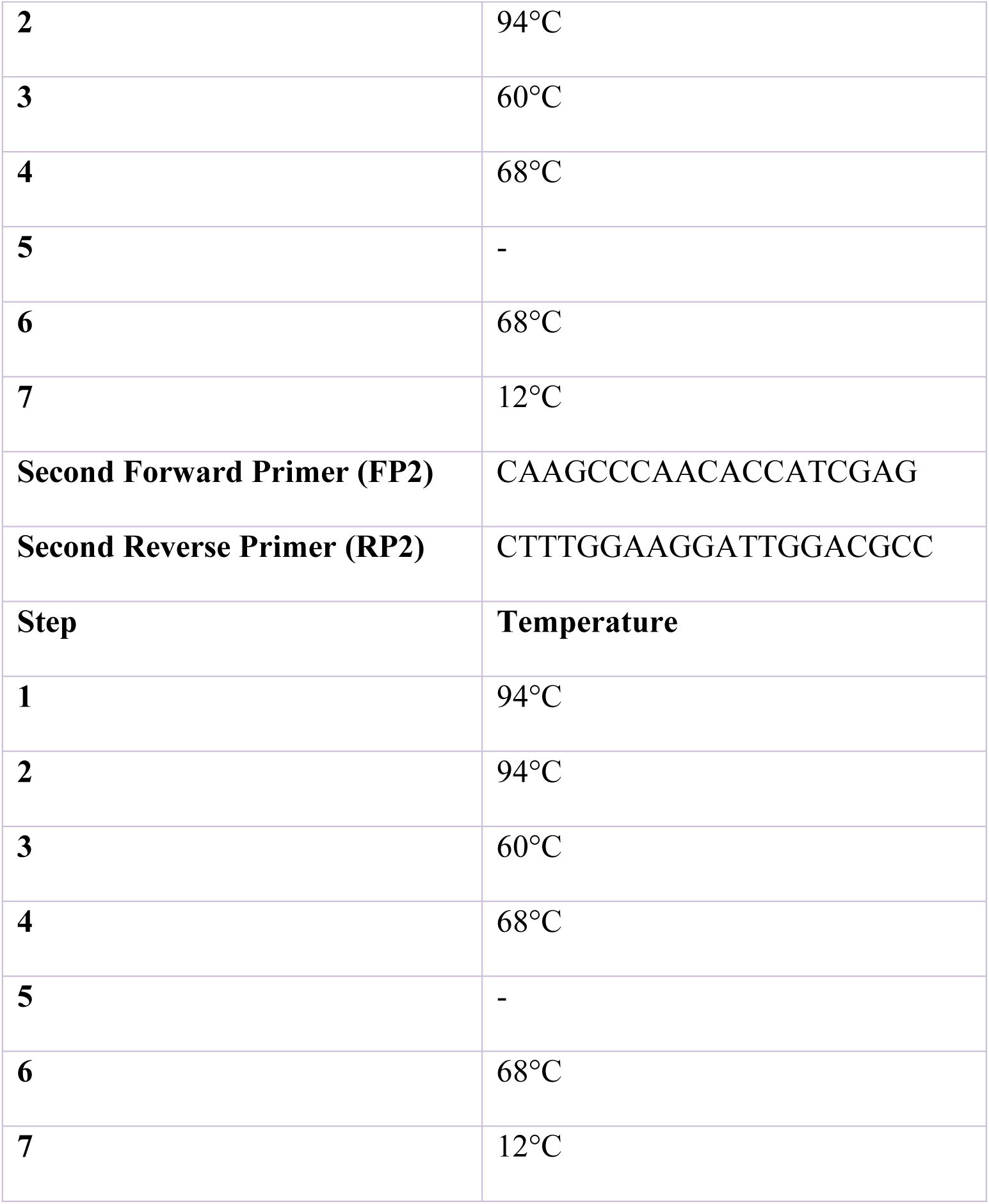

Adapter primers for Illumina sequencing were created and amplified by Azenta (‘Genewiz”) Inc. (Forward Primer: 5’-ACACTCTTTCCCTACACGACGCTCTTCCGATCT-3’; Reverse Primer: 5’-GACTGGAGTTCAGACGTGTGCTCTTCCGATCT-3’). All subsequent library preparations and Illumina 2×150 bp sequencing was performed by Azenta (‘Genewiz”) Inc. The samples were sequenced via alignment of fastq files and quantification of allele frequency for individual loci were performed using CRISPResso2 in batch mode^63^. The allele frequency for each site was calculated as the number of reads containing nucleotides C/G at the R1872W locus and the total number of variants reads containing nucleotides T/A at the R1872W locus in the application with default parameters with noise and low alignments to target sgRNA sequence filtered.

### Whole-Transcriptome RNA Sequencing

5 *Scn8a*^W/+^-EMX1 Sham and 5 *SCN8A*-ABE treated mice were taken down; hippocampal and cortical tissues were immediately dissected and collected for evaluation of RNA transcripts. Total RNA was harvested from cells in the hippocampus and cortex using the RNeasy Mini kit (Qiagen). Library preparation and sequencing were performed by Azenta “Genewiz”. Sequencing and subsequent FASTQs were performed and generated by Azenta “Genewiz” in paired-end mode on Illumina Novaseq. FastQC (v0.11.5) was used to assess the quality of each sample. Cutadapt (v3.4) was used to trim adapters. Reads were aligned to the mouse mm39 reference genome using STAR (v2.7.9a), and gene read counts were generated using Feature Counts (v2.0.6). Default parameters to remove low-quality bases, adapter sequences, reads under 3, and unpaired sequences. Trimmed reads were then aligned to the GENCODE mouse reference genomeGRCm39 using IGV genome analysis browser and refined to canonical coding sequences using CCDS release 21^64^ to confirm the absence of editing at all 27 sites identified for off-target effects within coding regions of DNA that are expressed in brain tissue. Expression of the mutant and wild-type alleles in heterozygous *Scn8a*^W/+^-EMX1 mice confirmed equal allelic expression of both mutant and ABE-treated groups. This was accomplished by detection of 2 (ATC-to-ATT) TALEN-induced silent mutations in exons 26b and 26a^34^ on the same platforms as described above.

### Whole-Genome Sequencing

2 *Scn8a*^W/+^-EMX1 Sham and 3 *SCN8A*-ABE treated mice were taken down; hippocampal and cortical tissues were immediately dissected and collected for evaluation of WGS. Total DNA was harvested from cells in the hippocampus and cortex using the Purelink Genomic DNA Mini Kit (Invitrogen). Library preparation and sequencing were performed by Azenta “Genewiz”. Sequencing and subsequent FASTQs were performed and generated by Azenta “Genewiz. QC controlled reads were then aligned to the GENCODE mouse reference genomeGRCm38 using IGV genome analysis browser to confirm the absence of editing at all 114 sites identified for off-target effects within mouse genomic DNA.

### Behavioral Assay: Open Field Test

All behavioral tests were conducted in behavioral testing rooms between 10:00 and 14:00 hours of the day during the light phase of the light/dark cycle. Behavioral tests were performed in mice between the ages of 4 and 8 weeks. Mice were tested in a random order. After the tests, the equipment was cleaned with 70% ethanol to eliminate olfactory cues. Exploratory behavior, anxiety-like behavior, and general locomotor activity were examined using the open-field test according to previous studies^65,66^. Each mouse was placed in the center of the apparatus consisting of a square area surrounded by white acrylic walls (45 × 45 × 40 cm). The total distance travelled (m) and time spent in the central area (s) were recorded. The central area was defined as the middle 20 × 20 cm area of the field. Data was collected over a 5-min period using the ANY-MAZE software. Data analysis was performed in Microsoft Excel. All statistical comparisons were made using the appropriate test in GraphPad Prism 10.

### Statistical Analysis

Survival analysis was compared by Mantel-Cox comparison test. Seizure incidence was compared by Mann-Whitney comparisons test. Comparisons of two groups, namely genomic data, were compared by an unpaired t-test when the data were normally distributed with equal variances, by Welch’s t-test when the data were normally distributed with unequal variances, and by Mann-Whitney test when the data were not normally distributed. Comparisons of three groups, namely, membrane and AP properties, peak sodium currents, and behavioral studies, were compared by one-way ANOVA followed by Dunnett’s multiple comparisons test when the data were normally distributed with equal variances, by Brown-Forsythe ANOVA with Dunnett’s multiple comparisons test when the data were normally distributed with unequal variances, and by the nonparametric Kruskal–Wallis test followed by Dunn’s multiple comparisons test when the data were not normally distributed. Data were assessed for normality using the Shapiro-Wilk test. Data were tested for outliers using the ROUT method to identify outliers. A two-way ANOVA followed by Tukey’s test for multiple comparisons was used to compare groups in experiments in which repetitive measures were made from a single cell over various voltage commands or current injections. Data are presented as individual data points and/or mean ± SEM. Exact n and P-values are reported in figure legends.

## Acknowledgements

This work was funded by National Institutes of Health (NIH) grants NS103090, NS122834, and NS120702 to MKP; NS34509 to MHM and F31 NS134264 to RMM. A Virginia Brain Institute Presidential Fellowship to CMR; a Virginia Brain Institute Transformative Neuroscience Pilot Grant to MKP and an Ivy Biomedical Innovation Fund Award to MKP; Special thanks to Brian Ruis, University of Virginia Genome Engineering Shared Resource Core; and Wenxi Yu, PhD for helpful discussions. Figures 1.a-b, 2.a-b, and 3.a were created with BioRender.com.

## Notes

### Competing Interest Statement

The authors have declared no competing interest.

### Summary of Updates

This version of the manuscript has been revised to include new off target validation and immunohistochemistry images.

## References

1. O’Brien, J. E. & Meisler, M. H. Sodium channel SCN8A (Nav1.6): properties and de novo mutations in epileptic encephalopathy and intellectual disability. Front. Genet. 4, 213 (2013).

2. Caldwell, J. H., Schaller, K. L., Lasher, R. S., Peles, E. & Levinson, S. R. Sodium channel Nav1.6 is localized at nodes of Ranvier, dendrites, and synapses. Proc. Natl. Acad. Sci. U. S. A. 97, 5616 (2000).

3. Veeramah, K. R. et al. De novo pathogenic SCN8A mutation identified by whole-genome sequencing of a family quartet affected by infantile epileptic encephalopathy and SUDEP. Am. J. Hum. Genet. 90, 502–510 (2012).

4. Meisler, M. SCN8A encephalopathy: Mechanisms and Models. Epilepsia 60, S86 (2019).

5. Talwar, D. & Hammer, M. F. *SCN8A* Epilepsy, Developmental Encephalopathy, and Related Disorders. Pediatr. Neurol. 122, 76–83 (2021).

6. Johannesen, K. M. et al. Early mortality in *SCN8A*-related epilepsies. Epilepsy Res. 143, 79–81 (2018).

7. Wagnon, J. L. et al. Pathogenic mechanism of recurrent mutations of SCN8A in epileptic encephalopathy. Ann. Clin. Transl. Neurol. 3, 114 (2015).

8. Magielski, J. H. et al. Deciphering the Natural History of SCN8A-Related Disorders. Neurology 104, e213533 (2025).

9. Fryxell, K. J. & Moon, W.-J. CpG mutation rates in the human genome are highly dependent on local GC content. Mol. Biol. Evol. 22, 650–658 (2005).

10. Bunton-Stasyshyn, R. K. A. et al. Prominent role of forebrain excitatory neurons in SCN8A encephalopathy. Brain 142, 362 (2019).

11. Ottolini, M., Barker, B. S., Gaykema, R. P., Meisler, M. H. & Patel, M. K. Aberrant Sodium Channel Currents and Hyperexcitability of Medial Entorhinal Cortex Neurons in a Mouse Model of SCN8A Encephalopathy. J. Neurosci. 37, 7643–7655 (2017).

12. Meisler, M. H. et al. SCN8A encephalopathy: Research progress and prospects. Epilepsia 57, 1027–1035 (2016).

13. Ademuwagun, I. A., Rotimi, S. O., Syrbe, S., Ajamma, Y. U. & Adebiyi, E. Voltage Gated Sodium Channel Genes in Epilepsy: Mutations, Functional Studies, and Treatment Dimensions. Front. Neurol. 12, 600050 (2021).

14. Møller, R. S. & Johannesen, K. M. Precision Medicine: SCN8A Encephalopathy Treated with Sodium Channel Blockers. Neurother. J. Am. Soc. Exp. Neurother. 13, 190–191 (2016).

15. Liu, D. R., Komor, A. C., Rees, H. A. & Kim, Y. Nucleobase editors and uses thereof. (2017).

16. Gaudelli, N. M. et al. Programmable base editing of A•T to G•C in genomic DNA without DNA cleavage. Nature 551, 464–471 (2017).

17. Richter, M. F., et al. Phage-assisted evolution of an adenine base editor with improved Cas domain compatibility and activity. Nat. Biotechnol. 38, 883–891 (2020).

18. Chatterjee, P. et al. A Cas9 with PAM recognition for adenine dinucleotides. Nat. Commun. 11, 2474 (2020).

19. Miller, S. M. et al. Continuous evolution of SpCas9 variants compatible with non-G PAMs. Nat. Biotechnol. 38, 471–481 (2020).

20. Walton, R. T., Christie, K. A., Whittaker, M. N. & Kleinstiver, B. P. Unconstrained genome targeting with near-PAMless engineered CRISPR-Cas9 variants. Science 368, 290–296 (2020).

21. Arbab, M. et al. Base editing rescue of spinal muscular atrophy in cells and in mice. Science 380, eadg6518 (2023).

22. Arbab, M. et al. Determinants of Base Editing Outcomes from Target Library Analysis and Machine Learning. Cell 182, 463–480.e30 (2020).

23. Concordet, J.-P. & Haeussler, M. CRISPOR: intuitive guide selection for CRISPR/Cas9 genome editing experiments and screens. Nucleic Acids Res. 46, W242 (2018).

24. Park, J., Bae, S. & Kim, J.-S. Cas-Designer: a web-based tool for choice of CRISPR-Cas9 target sites. Bioinforma. Oxf. Engl. 31, 4014–4016 (2015).

25. Zettler, J., Schütz, V. & Mootz, H. D. The naturally split Npu DnaE intein exhibits an extraordinarily high rate in the protein trans-splicing reaction. FEBS Lett. 583, 909–914 (2009).

26. Levy, J. M. et al. Cytosine and adenine base editing of the brain, liver, retina, heart and skeletal muscle of mice via adeno-associated viruses. *Nat*. Biomed. Eng. 4, 97 (2020).

27. Stoica, L., Ahmed, S. S., Gao, G. & Esteves, M. S. AAV-mediated gene transfer to the mouse CNS. Curr. Protoc. Microbiol. 29, 14D.5.1–14D.5.18 (2013).

28. Ryu, S.-M. et al. Adenine base editing in mouse embryos and an adult mouse model of Duchenne muscular dystrophy. Nat. Biotechnol. 36, 536–539 (2018).

29. Wenker, I. C. et al. Postictal Death Is Associated with Tonic Phase Apnea in a Mouse Model of Sudden Unexpected Death in Epilepsy. Ann. Neurol. 89, 1023–1035 (2021).

30. Deverman, B. E. et al. Cre-dependent selection yields AAV variants for widespread gene transfer to the adult brain. Nat. Biotechnol. 34, 204–209 (2016).

31. Chan, K. Y. et al. Engineered AAVs for efficient noninvasive gene delivery to the central and peripheral nervous systems. Nat. Neurosci. 20, 1172–1179 (2017).

32. Mathiesen, S. N., Lock, J. L., Schoderboeck, L., Abraham, W. C. & Hughes, S. M. CNS Transduction Benefits of AAV-PHP.eB over AAV9 Are Dependent on Administration Route and Mouse Strain. Mol. Ther. Methods Clin. Dev. 19, 447 (2020).

33. Cradick, T. J., Qiu, P., Lee, C. M., Fine, E. J. & Bao, G. COSMID: A Web-based Tool for Identifying and Validating CRISPR/Cas Off-target Sites. Mol. Ther. Nucleic Acids 3, e214 (2014).

34. Jones, J. M. & Meisler, M. H. Modeling human epilepsy by TALEN targeting of mouse sodium channel Scn8a. Genesis 52, 141–148 (2014).

35. Wengert, E. R. & Patel, M. K. The Role of the Persistent Sodium Current in Epilepsy. Epilepsy Curr. 21, 40–47 (2020).

36. Kraeuter, A.-K., Guest, P. C. & Sarnyai, Z. The Open Field Test for Measuring Locomotor Activity and Anxiety-Like Behavior. Methods Mol. Biol. Clifton NJ 1916, 99–103 (2019).

37. Schaller, K. L., Krzemien, D. M., Yarowsky, P. J., Krueger, B. K. & Caldwell, J. H. A novel, abundant sodium channel expressed in neurons and glia. J. Neurosci. 15, 3231 (1995).

38. Han, Z. et al. Antisense oligonucleotides increase Scn1a expression and reduce seizures and SUDEP incidence in a mouse model of Dravet syndrome. Sci. Transl. Med. 12, eaaz6100 (2020).

39. Lenk, G. M. et al. Scn8a Antisense Oligonucleotide Is Protective in Mouse Models of SCN8A Encephalopathy and Dravet Syndrome. Ann. Neurol. 87, 339–346 (2020).

40. Hill, S. F., Ziobro, J. M., Jafar-Nejad, P., Rigo, F. & Meisler, M. H. Genetic interaction between Scn8a and potassium channel genes Kcna1 and Kcnq2. Epilepsia 63, e125–e131 (2022).

41. Hill*, S. F., et al. Long-Term Downregulation of the Sodium Channel Gene Scn8a Is Therapeutic in Mouse Models of Epilepsy. Ann. Neurol. 95, 754–759 (2024).

42. Yuan, Y. et al. Antisense oligonucleotides restore excitability, GABA signalling and sodium current density in a Dravet syndrome model. Brain J. Neurol. 147, 1231–1246 (2024).

43. Cantara, S., Simoncelli, G. & Ricci, C. Antisense Oligonucleotides (ASOs) in Motor Neuron Diseases: A Road to Cure in Light and Shade. Int. J. Mol. Sci. 25, 4809 (2024).

44. Crooke, S. T. Antisense Drug Technology: Principles, Strategies, and Applications, Second Edition. (Taylor & Francis, 2007).

45. Yu, W. et al. Allele-Specific Editing of a Dominant Epilepsy Variant Protects against Seizures and Lethality in a Murine Model. Ann. Neurol. 96, 958–969 (2024).

46. Wong, J. C. et al. Autistic-like behavior, spontaneous seizures, and increased neuronal excitability in a Scn8a mouse model. Neuropsychopharmacology 46, 2011–2020 (2021).

47. Conecker, G. et al. Global modified-Delphi consensus on comorbidities and prognosis of SCN8A-related epilepsy and/or neurodevelopmental disorders. Epilepsia 65, 2308–2321 (2024).

48. Baker, E. M. et al. The novel sodium channel modulator GS-458967 (GS967) is an effective treatment in a mouse model of SCN8A encephalopathy. Epilepsia 59, 1166–1176 (2018).

49. Johnson, J. et al. NBI-921352, a first-in-class, NaV1.6 selective, sodium channel inhibitor that prevents seizures in Scn8a gain-of-function mice, and wild-type mice and rats. eLife 11, e72468 (2022).

50. Wu, L., et al. Adenine base editors induce off-target structure variations in mouse embryos and primary human T cells. Genome Biol. 25, 291 (2024).

51. Doench, J. G. et al. Optimized sgRNA design to maximize activity and minimize off-target effects of CRISPR-Cas9. Nat. Biotechnol. 34, 184–191 (2016).

52. Gardella, E. & Møller, R. S. Phenotypic and genetic spectrum of SCN8A-related disorders, treatment options, and outcomes. Epilepsia 60 Suppl 3, S77–S85 (2019).

53. Wagnon, J. L. & Meisler, M. H. Recurrent and Non-Recurrent Mutations of SCN8A in Epileptic Encephalopathy. Front. Neurol. 6, 104 (2015).

54. Matsuda, T. & Cepko, C. L. Controlled expression of transgenes introduced by in vivo electroporation. Proc. Natl. Acad. Sci. U. S. A. 104, 1027–1032 (2007).

55. Herzog, R. I., Cummins, T. R., Ghassemi, F., Dib-Hajj, S. D. & Waxman, S. G. Distinct repriming and closed-state inactivation kinetics of Nav1.6 and Nav1.7 sodium channels in mouse spinal sensory neurons. J. Physiol. 551, 741–750 (2003).

56. Kluesner, M. G. et al. EditR: A Method to Quantify Base Editing from Sanger Sequencing. CRISPR J. 1, 239–250 (2018).

57. Kobayashi, Y. & Hensch, T. Germline recombination by conditional gene targeting with Parvalbumin-Cre lines. Front. Neural Circuits 7, 168 (2013).

58. Luo, L. et al. Optimizing Nervous System-Specific Gene Targeting with Cre Driver Lines: Prevalence of Germline Recombination and Influencing Factors. Neuron 106, 37–65.e5 (2020).

59. Wengert, E. R. et al. Targeted Augmentation of Nuclear Gene Output (TANGO) of *Scn1a* rescues parvalbumin interneuron excitability and reduces seizures in a mouse model of Dravet Syndrome. Brain Res. 1775, 147743 (2022).

60. Koblan, L. W. et al. In vivo base editing rescues Hutchinson–Gilford progeria syndrome in mice. Nature 589, 608–614 (2021).

61. Hargus, N. J. et al. Temporal lobe epilepsy induces intrinsic alterations in Na channel gating in layer II medial entorhinal cortex neurons. Neurobiol. Dis. 41, 361–376 (2011).

62. Wenker, I. C., et al. Forebrain epileptiform activity is not required for seizure-induced apnea in a mouse model of Scn8a epilepsy. Front. Neural Circuits 16, (2022).

63. Clement, K. et al. CRISPResso2 provides accurate and rapid genome editing sequence analysis. Nat. Biotechnol. 37, 224–226 (2019).

64. Pujar, S. et al. Consensus coding sequence (CCDS) database: a standardized set of human and mouse protein-coding regions supported by expert curation. Nucleic Acids Res. 46, D221–D228 (2018).

65. Ueno, H. et al. Effects of repetitive gentle handling of male C57BL/6NCrl mice on comparative behavioural test results. Sci. Rep. 10, 3509 (2020).

66. Umemura, M. et al. Comprehensive Behavioral Analysis of Activating Transcription Factor 5-Deficient Mice. Front. Behav. Neurosci. 11, 125 (2017).

