## Supplemental Tables & Figures for "Base Editing Rescue of Seizures and SUDEP in *SCN8A* Developmental Epileptic Encephalopathy"

Supplemental Information

| Construst Screen Name | 16 R1872W Targeting Guide RNA Sequences | PAM | Base Editor and Construct Type |
|---|---|---|---|
| C16 | GCACGAACCaCTCCTCCATCTGCT | GCCGCA | eNme2-C-ABE8e.v4 |
| C15 | GCGCCACGAACCaCTCCTCCATCT | CTGCCG | eNme2-C-ABE8e.v3 |
| C14 | GCCACGAACCaCTCCTCCATCTG | GCTGCC | eNme2-C-ABE8e.v2 |
| C13 | GACGCCACGAACCaCTCCTCCA | TCTGCT | eNme2-C-ABE8e.v1 |
| C12 | GCGAACCaCTCCTCCATCTG | CTG | ABE8e-SpRY.v5 |
| C11 | GACCaCTCCTCCATCTGCTGCC | GCA | ABE8e-SpRY.v4 |
| C10 | GAACCaCTCCTCCATCTGCT | GCC | ABE8e-SpRY.v2 |
| C9 | GAACCaCTCCTCCATCTGCTG | CCG | ABE8e-SpRY.v1 |
| C8 | GGAACCaCTCCTCCATCTGC | TGCC | ABEmax-NRCH.v4 |
| C7 | GCGAACCaCTCCTCCATCTGC | TGCC | ABEmax-NRCH.v3 |
| C6 | GACCaCTCCTCCATCTGCTGCC | CGCA | ABEmax-NRCH.v2 |
| C5 | GAACCaCTCCTCCATCTGCTGC | CGCA | ABEmax-NRCH.v1 |
| C4 | GGAACCaCTCCTCCATCTGC | TGCC | ABE8e-NRCH.v4 |
| C3 | GCGAACCaCTCCTCCATCTGC | TGCC | ABE8e-NRCH.v3 |
| C2 | GACCaCTCCTCCATCTGCTGCC | CGCA | ABE8e-NRCH.v2 |
| C1 | GAACCaCTCCTCCATCTGCTGC | CGCA | ABE8e-NRCH.v1 |

**Supplementary Table 1: R1872W targeting guide RNA sequences and ABEs for cell screen assays.**
A panel of the 16 chosen guide RNA (gRNA) sequences paired with a specific base editor construct targeting the R1872W mutation was designed for base editing. Each gRNA sequence is associated with a PAM motif.

| Construct Name | Base Editor and Construct Type | Position of (A) in sgRNA | PAM |
|---|---|---|---|
| C16 | eNme2-C-ABE8e.v4 | 9 | GCCGCA |
| C14 | eNme2-C-ABE8e.v2 | 11 | CTGCCG |
| C15 | eNme2-C-ABE8e.v3 | 12 | GCTGCC |
| C13 | eNme2-C-ABE8e.v1 | 15 | TCTGCT |
| C12 | ABE8e-SpRY.v5 | 8 | CTG |
| C11 | ABE8e-SpRY.v4 | 5 | GCA |
| C10 | ABE8e-SpRY.v2 | 6 | GCC |
| C9 | ABE8e-SpRY.v1 | 6 | CCG |
| C8 | ABEmax-NRCH.v4 | 7 | TGCC |
| C7 | ABEmax-NRCH.v3 | 8 | TGCC |
| C6 | ABEmax-NRCH.v2 | 5 | CGCA |
| C5 | ABEmax-NRCH.v1 | 6 | CGCA |
| C4 | ABE8e-NRCH.v4 | 7 | TGCC |
| C3 | ABE8e-NRCH.v3 | 8 | TGCC |
| C2 | ABE8e-NRCH.v2 | 5 | CGCA |
| C1 | ABE8e-NRCH.v1 | 6 | CGCA |
|  | HEK293 Control |  |  |

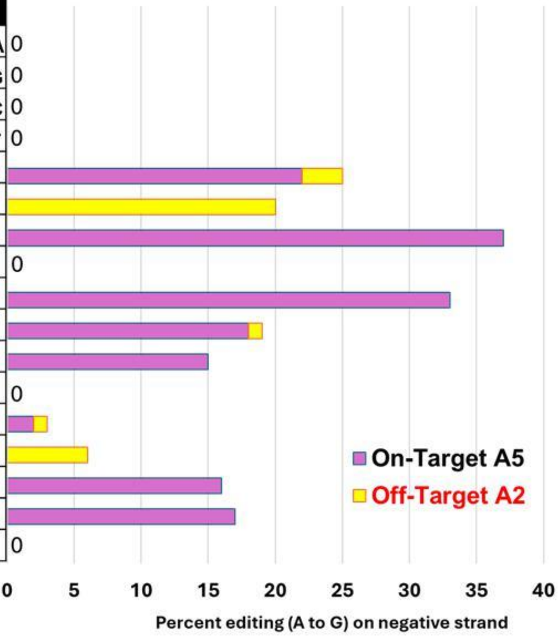


**Supplementary Figure 1. ABE efficiencies of 16 constructs in R1872W expressing HEK293- cells**.
Results of Sanger sequencing for on-target (A5, pink) and off-target (A2, yellow) A-to-G editing efficiencies for base editor constructs within the targeted sequence. X-axis shows percent editing on the negative strand of the R1872W locus. Analysis of files were conducted with EditR1.0.10 with default parameters.

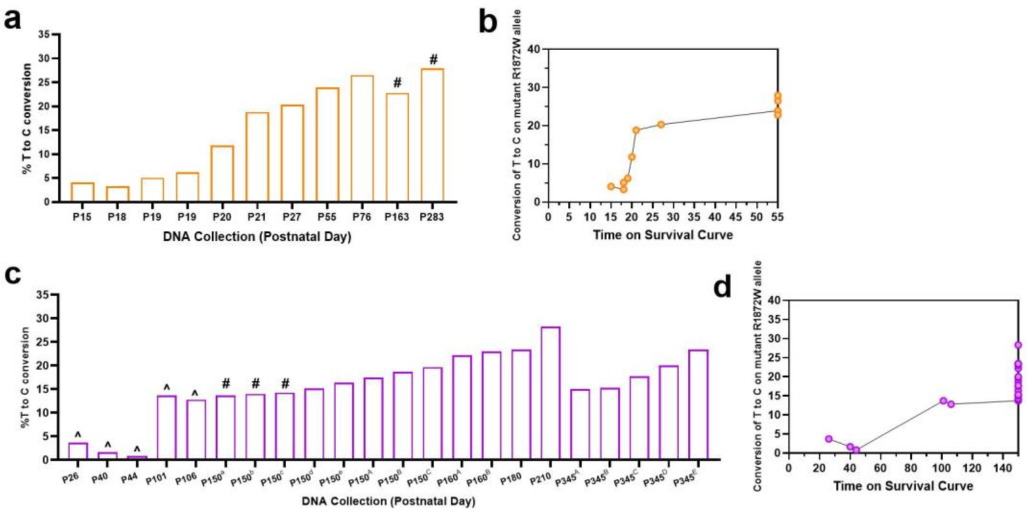


e

| ABE Significant Survival DNA Base Ratios | DNA NGS | DNA NGS |
|---|---|---|
| Base Percentage at R1872W | Wildtype | Mutant |
| *Scn8a*W/+-Ella Sham | 6.9% | 93.1% |
| *Scn8a*W/+-Ella ABE | 29.1% | 70.9% |
| ABE Treatment Editing Results | 22.2% | -22.2% |

f

| ABE Significant Survival DNA Base and Transcript Ratio | DNA NGS | DNA NGS | RNA-Seq | RNA-Seq |
|---|---|---|---|---|
| Base Percentage at R1872W | Wildtype | Mutant | Wildtype | Mutant |
| *Scn8a*W/+-EMX1 Sham | 26.3% | 73.7% | 56.6% | 43.4% |
| *Scn8a*W/+-EMX1 ABE | 44.5% | 55.5% | 90.0% | 10.0% |
| ABE Treatment Editing Results | 18.2% | -18.2% | 33.4% | -33.4% |

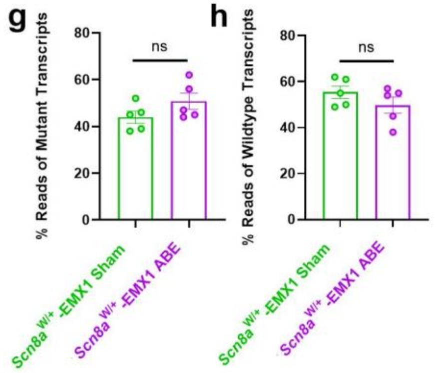


**Supplementary Figure 2. ABE-mediated editing and survival outcomes**

(**a**) Percentages of each mouse of ABE-mediated T-to-C conversion on the mutant *SCN8A* R1872W allele in *Scn8a* $^{W/+}$-EIIa ABE treated mice (n=11) at different survival time points. # denotes mice that were euthanized for sequencing purposes. No symbol above the bar indicates mice that were found dead immediately before sample was collected. (**b**) Longitudinal analysis of T-to-C conversion at the R1872W allele plotted against survival time in *Scn8a*$^{W/+}$-EIIa ABE (n=11) mice. Each point represents an individual mouse. (**c**) Percentage of T-to-C conversion on the mutant R1872W allele in *Scn8a* $^{W/+}$-EMX1 ABE-treated mice (n=22) at different survival time points. ^ denotes mice that experienced premature death. # denotes mice that were included in the EEG evaluations and experienced seizures but no premature death. No symbol above the bar indicates mice that were euthanized for sequencing purposes. Each point represents an individual mouse. (**d**) Longitudinal analysis of T-to-C conversion at the R1872W allele plotted against survival time in *Scn8a*$^{W/+}$-EMX1 ABE (n=22) mice. Each point represents an individual mouse. (**e**) Average genomic reads quantification of *Scn8a*$^{W/+}$-EIIa Sham mice (n=9) compared to ABE-treated *Scn8a*$^{W/+}$-EIIa mice (n=6) that survived past P21, a timepoint where all *Scn8a*$^{W/+}$-EIIa Sham mice had succumbed to death. (**f**) Average genomic reads quantification of *Scn8a*$^{W/+}$-EMX1 Sham mice (n=19) compared to ABE-treated *Scn8a*$^{W/+}$-EMX1 mice (n=19) that survived past P65, a timepoint where all *Scn8a*$^{W/+}$-EMX1 Sham mice had succumbed to death. (**g**) Expression of the mutant alleles in heterozygous *Scn8a*$^{W/+}$-EMX1 mice confirmed equal allelic expression of both mutant (n=5) and ABE-treated (n=5) groups. (**h**) Expression of the wild-type alleles in heterozygous *Scn8a*$^{W/+}$-EMX1 mice confirmed equal allelic expression of both mutant (n=5) and ABE-treated(n=5) groups. Data represents mean ± S.E.M.

### Supplementary Table 2: Membrane, AP, and channel properties of *Scn8a*$^{W/+}$-EMX1 neurons

| | $V_m$ (mV) | AP threshold (mV) | Rheobase (pA) | Upstroke Velocity (mV/ms) | Downstroke Velocity (mV/ms) | Amplitude (mV) | $APD_{50}$ (ms) | Input Resistance (MΩ) | $V_{1/2}$ (mV) |
|---|---|---|---|---|---|---|---|---|---|
| *Scn8a*$^{W/+}$-control (*n*=21, 4; *n*=17, 6) | -67.0 ± 1.0 | -46.7 ± 0.8 | 126.7 ± 14.1 | 283.4 ± 13.9 | -64.7 ± 6.9 | 91.5 ± 2.5 | 1.4 ± 0.1 | 101.8 ± 8.8 | -58.0 ± 2.4 |
| *Scn8a*$^{W/+}$-EMX1 ABE (*n*=34, 7; *n*=17, 8) | -67.4 ± 0.9 | -45.0 ± 0.6 | 173.5 ± 12.3 * | 389.3 ± 20.8 *** | -122.0 ± 6.9 **** | 83.2 ± 1.8 * | 0.84 ± 0.0 **** | 76.9 ± 4.6* | -61.0 ± 1.7 |
| *Scn8a*$^{W/+}$-EMX1 Sham (*n*=25, 4; *n*=24, 7) | -66.1 ± 0.9 | -46.0 ± 0.7 | 152.0 ± 10.7 | 382.3 ± 23.4 | -98.0 ± 5.5 **, # | 88.0 ± 1.7 | 1.0 ± 0.1 ** | 81.7 ± 5.5 | -61.6 ± 1.7 |

Recordings were carried out in pyramidal neurons from Layer IV/V somatosensory cortex of *Scn8a*$^{W/+}$-control, *Scn8a*$^{W/+}$-EMX1 ABE, and *Scn8a*$^{W/+}$-EMX1 Sham mice at P17-25. Recordings were performed in multiple cells from each animal (*n*=cells, animals) and in multiple experiments as shown here: genotype (membrane/AP properties *n*; channel properties *n*).

*, $P<0.05$; **, $P<0.01$; ***, $P<0.001$; ****, $P<0.0001$ indicate significance vs. *Scn8a*$^{W/+}$-control mice.

#, $P<0.05$; ##, $P<0.01$; ####, $P<0.0001$ indicate significance vs. *Scn8a*$^{W/+}$-EMX1 ABE-treated mice. Data are presented as mean ± SEM.

### Supplementary Table 3: Membrane and AP properties of *Scn8a*$^{W/+}$-EIIa neurons

| | $V_m$ (mV) | AP threshold (mV) | Rheobase (pA) | Upstroke Velocity (mV/ms) | Downstroke Velocity (mV/ms) | Amplitude (mV) | $APD_{50}$ (ms) | Input Resistance (MΩ) |
|---|---|---|---|---|---|---|---|---|
| *Scn8a*$^{W/+}$-control (n=29,5) | -64.7 ± 0.6 | -43.9 ± 0.6 | 81.4 ± 6.9 | 269.5 ± 14.8 | -62.0 ± 3.2 | 84.4 ± 2.3 | 1.4 ± 0.1 | 120.5 ± 5.8 |
| *Scn8a*$^{W/+}$-EIIa ABE (*n*=26, 8) | -67.4 ± 1.1 | -46.7 ± 0.9 # | 111.5 ± 9.4 * | 377.2 ± 20.3 *** | -69.8 ± 4.4 | 86.9 ± 1.7 | 1.2 ± 0.1 | 110.4 ± 7.9 |
| *Scn8a*$^{W/+}$-EIIa Sham (*n*=17, 5) | -64.2 ± 0.9 | -47.6 ± 1.5 * | 72.9 ± 11.2 # | 285.9 ± 17.0 # | -58.2 ± 5.0 | 89.1 ± 2.1 | 1.6 ± 0.1 ## | 136.7 ± 11.0 |

Recordings were carried out in pyramidal neurons from Layer IV/V somatosensory cortex of *Scn8a*$^{W/+}$-control, *Scn8a*$^{W/+}$-EIIa ABE, and *Scn8a*$^{W/+}$-EIIa Sham mice at P13-17. Recordings were performed in multiple cells from each animal (n=cells, animals). Data are presented as mean ± SEM.

*, $P<0.05$; ***, $P<0.001$ indicate significance when compared to *Scn8a*$^{W/+}$-control mice.

#, $P<0.05$; ##, $P<0.01$; ###, $P<0.001$ indicate significance when compared to *Scn8a*$^{W/+}$-EIIa ABE mice.

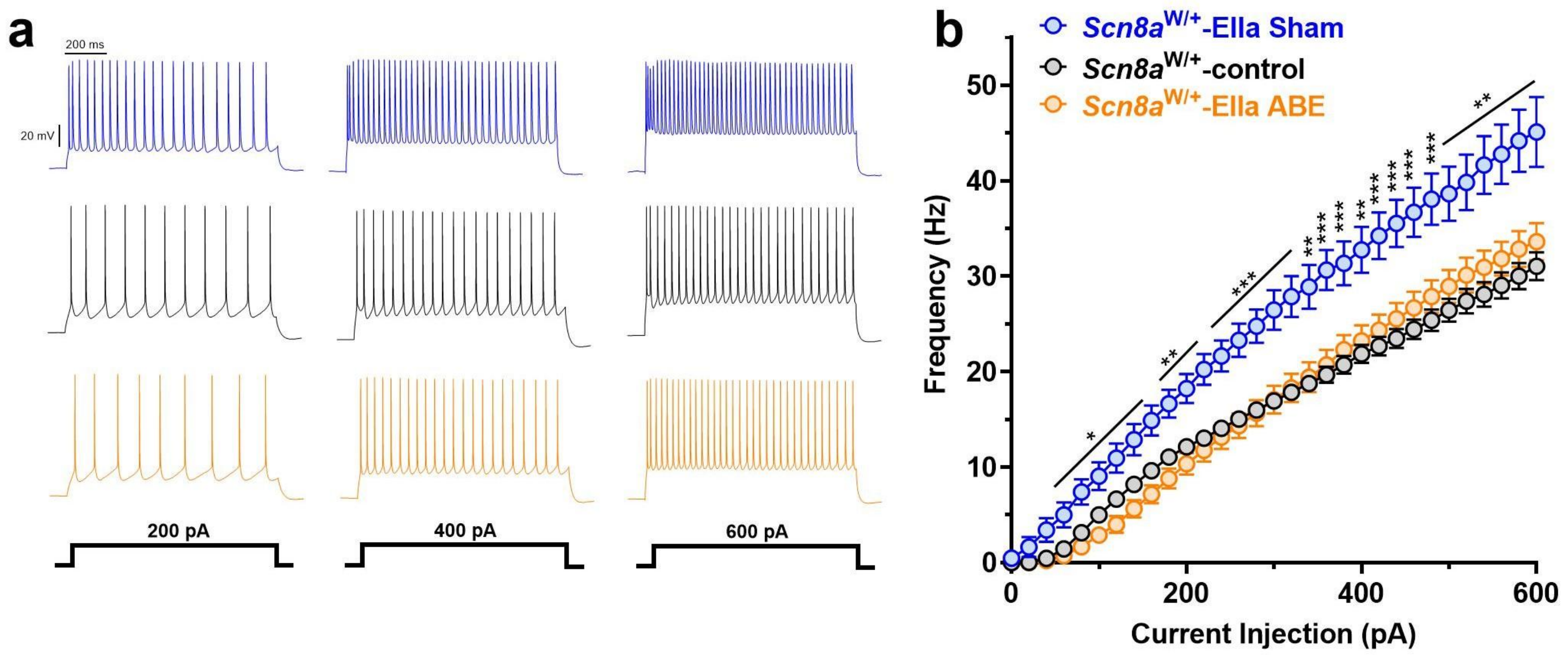


**Supplementary Figure 3. *SCN8A*-ABE attenuates neuronal hyperexcitability in *Scn8a*$^{W/+}$-EIIa mice.**
(**a**) Example traces of action potential firing from *Scn8a*$^{W/+}$-EIIa Sham (blue), *Scn8a*$^{W/+}$-control (black), and *Scn8a*$^{W/+}$-EIIa ABE (orange) neurons at current injection steps of 200 pA, 400 pA, or 600 pA. (**b**) *Scn8a*$^{W/+}$-EIIa Sham (n=17, 5 mice) neurons exhibit increased firing when compared to *Scn8a*$^{W/+}$-control (n=29, 5 mice) that is attenuated in *Scn8a*$^{W/+}$-EIIa ABE neurons (n=26 cells, 8 mice; *, P<0.05; **, P<0.01; ***, P<0.001; ****, P<0.0001; two-way ANOVA with Dunns's multiple comparisons test).
